# Cytoplasmic poly-adenosine binding proteins modulate susceptibility of mRNAs to RNA-binding protein-directed decay

**DOI:** 10.1101/2025.10.02.680050

**Authors:** Katherine M. McKenney, Carmen Hernandez-Perez, Elise B. Dunshee, John M. Pum, Aaron C. Goldstrohm

## Abstract

The cytoplasmic fate of mRNAs is dictated by the balance of translation and mRNA degradation, governed in part by the 3′ poly-adenosine tail and cytoplasmic poly(A)-binding proteins (PABPCs). Deadenylases remove poly(A) to initiate mRNA decay, while sequence-specific RNA-binding factors, including Pumilio proteins (PUM1 and PUM2), modulate these processes. We investigated how human PUM1&2 repress target mRNAs by accelerating their degradation. We found that the poly(A) tail plays a central role in PUM repression, dependent on the interplay of deadenylases and PABPCs. PUM-mediated repression requires the CCR4-NOT deadenylase but not the poly(A) nuclease (PAN). PUMs associate with and require PABPC1 and PABPC4 to repress. In the absence of PABPCs, both PUM targets and non-targets become unstable, bypassing PUM control. Increasing PABPC inhibits PUM activity in a concentration-dependent manner by stabilizing poly(A) mRNAs. Our results establish a Goldilocks principle wherein PABPC abundance tunes the response of mRNAs to regulatory factors through protection of poly(A) from deadenylation. Variation of PABPC levels across tissues and development suggests physiological relevance for this mechanism.

## INTRODUCTION

Eukaryotic mRNAs are tightly controlled in the cytoplasm to establish the proper amount, timing, and location of protein expression. Regulation of translation and mRNA stability is governed by the unique modifications at each end of the transcript: the 5′ 7-methyl-guanosine “cap” structure and the 3′ poly-adenosine (poly(A)) tail. In the cytoplasm, these features are recognized by specific RNA-binding factors. The 5′ cap is bound by the translation initiation factor eIF4E. The poly(A) tail is bound by cytoplasmic poly(A)-binding proteins (PABPCs) that enhance translation and mRNA stability (1–4) by interacting with translation initiation factor eIF4G, which in turn binds to 5′ cap-bound eIF4E (5, 6).

PABPC is highly conserved throughout eukarya and humans encode five PABPC isoforms; PABPC1 is generally the most abundant (7). The N-terminal four RNA recognition motifs (RRMs) of PABPC1 bind to poly(A) with high affinity, a proline-rich linker supports cooperative assembly on poly(A), and a C-terminal MLLE motif engages with PAM2 containing protein partners. Additional human PABPC family members include the testes-specific PABPC3 (tPABP), inducible PABPC4 (iPABP), embryonic PABPC1L (ePABP), and ovary-enriched, X-linked PABPC5 (8–13). The functional similarities and differences among these PABPC paralogs remain incompletely defined.

Messenger RNA decay pathways act on the poly(A) tail and 5′ cap to initiate degradation. Shortening of the poly(A) tail by deadenylase enzymes initiates decay of most mRNAs (14–16). Multiple deadenylase enzyme complexes can act on poly(A) mRNAs, including the CCR4-NOT complex and the poly(A) nuclease (PAN) (14–16). Subsequently, the mRNA is rapidly degraded by two pathways. In 5′ decay, decapping precedes 5′→3′ degradation by XRN1 (14–16). The deadenylated mRNA can also be degraded in a 3′→5′ direction by the exosome complex (14–16).

The interplay of poly(A) and PABPC with deadenylases plays a central role in modulating mRNA stability (15). How PABPC protection of poly(A) is overcome in cells remains unsettled. In vitro, PABPC can block degradation of poly(A) (19–25). For PAN deadenylase, PABPC facilitates substrate recognition and stimulates shortening of long poly(A) tails (17, 18). In the case of CCR4-NOT, biochemical analysis indicated that its two deadenylase subunits differ in their sensitivity to PABPC1 (26, 27). In cells, analysis of deadenylation of reporter mRNAs indicated a sequential biphasic process whereby PAN first shortens the poly(A) tail and then CCR4-NOT completes deadenylation (28). However, a subsequent transcriptome-wide analysis supports a more restricted role for PAN in trimming long poly(A) tails while CCR4-NOT performs bulk deadenylation (26).

Several limitations hindered analysis of PABPC function in vivo. First, PABPC is essential for viability (29). Second, PABPC1 protein is abundant and often potentially redundant paralogs are co-expressed (29, 30). RNAi studies suggested that PABPC1 promotes deadenylation in HeLa cells (26); however, the extended RNAi-mediated depletion and co-expression of PABPC4 limit interpretation. Recently, degron-based systems have enabled rapid protein depletion, revealing the role of PABPC1 and PABPC4 in stabilizing mRNAs in vivo (29, 30).

Translation and decay rates of mRNAs vary widely, dictated by *cis*-acting RNA sequence elements and *trans*-acting RNA-binding proteins (RBPs) (14, 16, 31–33). Each mRNA is likely to be controlled by multiple factors that establish the proper level of translation and stability (14, 15, 34). A growing number of sequence specific RBPs have been shown to regulate their target mRNAs by modulating mRNA deadenylation (14, 31). A prime example is the Pumilio family.

Humans encode two paralogs, PUM1 and PUM2, that exhibit the same RNA-binding and repressive activities (33, 35–37). PUMs have conserved functions in development, fertility, neurological processes, and stem cell fate (33), and their dysfunction contributes to neuro- developmental disorders, epilepsy, and cancer (38–41). PUMs are defined by a unique RNA-binding domain that binds with high affinity to the specific eight nucleotide Pumilio Response Element (PRE) with the consensus 5′-UGUANAUA (N=U,A,G,C). PUMs repress PRE-containing target mRNAs by directly recruiting the CCR4-NOT deadenylase complex to accelerate their degradation (35, 37).

In this study, we analyzed the roles of poly(A), deadenylases, and PABPCs in the mechanism of repression by human PUM1&2. We found that the poly(A) tail of target mRNAs and the CCR4- NOT deadenylase are required for PUM repression, whereas PAN is not. PUMs physically associate with the PABPC1 and PABPC4 proteins, and we showed that both are necessary for repression of mRNAs by PUM. In fact, depletion of PABPCs destabilized both PUM target and non-target mRNAs, bypassing PUM-accelerated deadenylation. In contrast, increasing PABPC1 concentration in cells blocked repression and mRNA degradation by PUMs in a dose-dependent manner. This effect requires the RNA-binding activity of PABPC1, but not its ability to bind to protein partners. The data support a model wherein PABPCs primarily modulate the ability of PUMs to promote degradation of target mRNAs by protecting the poly(A) tail. The results emphasize the balance of poly(A):PABPC dependent stabilization and CCR4-NOT mediated deadenylation, illuminating how PUMs shift that balance toward decay. More broadly, our results illuminate how PABPC levels can tune responsiveness to RBP directed deadenylation and decay. Low levels of PABPC destabilize poly(A) mRNAs such that recruitment by CCR4- NOT by sequence specific RBPs is obviated. In contrast, high levels of PABPC stabilize mRNAs, making them resistant to decay and RBP-mediated deadenylation. Normal PABPC levels enable responsiveness to sequence-specific regulation. We provide evidence that PABPC levels vary widely across tissues, suggesting a broad physiological relevance to these results.

## RESULTS

### The poly(A) tail is required for repression by human Pumilio proteins

We previously showed that PUM1&2 directly recruit the CCR4-NOT deadenylase complex to promote decay of PRE-containing mRNAs (35, 37). If this is the major mechanism of PUM-mediated repression, then the poly(A) tail of the target mRNA should be necessary for PUM activity. To test this hypothesis, we used an established, cell-based, PUM reporter gene assay (35–37) wherein Nano-luciferase (Nluc) mRNA is controlled by three PREs in the context of a minimal 3′ UTR (**Figure 1A**, Nluc 3xPRE). Repression of Nluc 3xPRE by endogenous PUM1&2 is calculated relative to a mutant version wherein the 5′-UGU trinucleotide of each PRE, essential for PUM-binding, was converted to 5′-ACA (**Figure 1A**, Nluc 3xPREmt). A firefly luciferase (Fluc) expression plasmid was cotransfected to normalize for transfection efficiency. In the HCT116 human colorectal carcinoma cell line, endogenous PUM1&2 repressed the expression of the 3xPRE reporter relative to the 3xPREmt by 2.8-fold **(Figure 1B),** consistent with previous reports (35–37).

**Figure 1.**
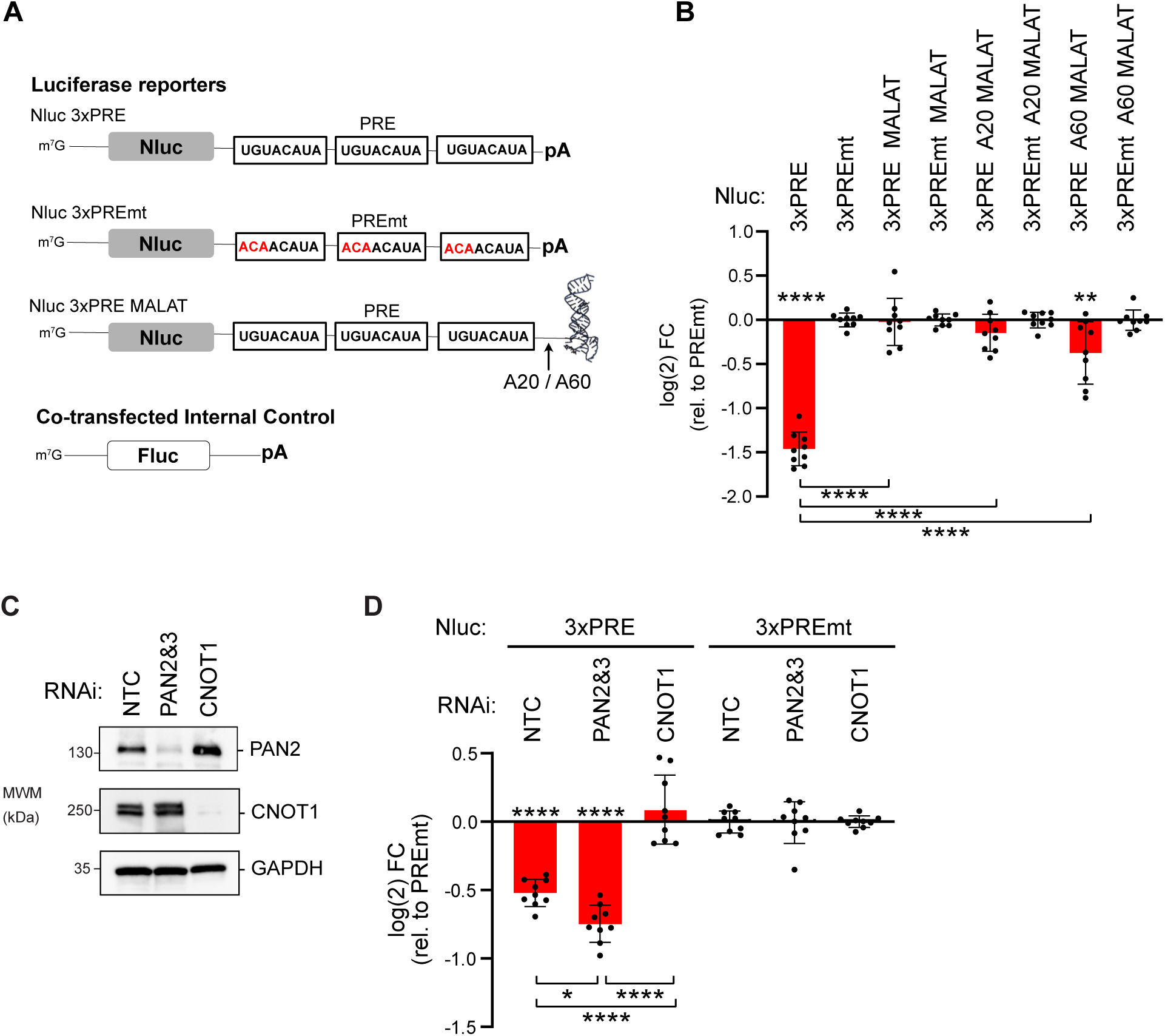
The poly(A) tail is necessary for PUM-mediated repression. **A.** PUM1&2 repression activity was measured in HCT116 cells using Nano-luciferase reporters with three PREs within a minimal 3′ UTR with cleavage and poly-adenylation signals (Nluc 3xPRE), calculated relative to a version wherein the PRE sequences were mutated (indicated in red text) to prevent PUM binding (Nluc 3xPREmt). PUM repression of the poly-adenylated reporter was compared to a Nluc reporter that has a 3′ end generated by the MALAT1 non-coding RNA (Nluc 3xPRE MALAT), which is processed by RNase P mediated cleavage to form a triple helix structure (PBD: 4PLX). Derivatives of the Nluc 3xPRE were constructed with internal poly(A) tracts of either 20 (A20) or 60 (A60) adenosines, inserted between the PREs and the MALAT1 triple helix. Firefly luciferase (Fluc) served as an internal control to normalize transfection efficiency. **B.** PRE dependent repression by endogenous PUM1 and PUM2 was measured as log(2) fold change (FC) of each Nluc 3xPRE reporter relative to its corresponding 3xPREmt reporter. Mean fold change is plotted along with individual replicate data points. n=9; 3 experiments, each with 3 biological replicates; +/- standard deviation (SD). For significance calling, p < 0.05 = *, p < 0.01 = **, p < 0.001 = ***, p < 0.0001 = **** based on ordinary one-way ANOVA and Tukey test for multiple comparisons. Asterisks above the axis denote significance relative to the 3xPREmt version of each reporter type, whereas below the bars are calculated relative to the poly-adenylated Nluc reporters. **C.** Western blot confirming the depletion of PAN2 and CNOT1 proteins by RNAi in HCT116 cells. GAPDH served as a loading control. n=3 experimental replicates. **D.** The effect of CCR4-NOT or PAN2 and PAN3 knockdown on PUM repression of the Nluc 3xPRE reporter, relative to the mutant Nluc 3xPREmt, was measured in HCT116 cells, in comparison to cells transfected with non-targeting control siRNAs. n=9; 3 experiments, each with 3 biological replicates; +/- SD.

We tested the requirement of the 3′ poly(A) tail by comparing PUM repression of the poly- adenylated Nluc 3xPRE to an identical mRNA that has a 3′ end terminating in the MALAT1 triple helix structure **(Figure 1A**, Nluc 3xPRE MALAT). Notably, the MALAT1 3′ end was previously reported to stabilize and support translation of an mRNA (42–44). Consistent with this, our Nluc MALAT reporters were expressed at similar levels to their poly(A) counterparts (**Figure S1**). We found that PUMs did not repress the Nluc 3xPRE MALAT relative to the version with mutant PREs, Nluc 3xPREmt MALAT **(Figure 1B)**, indicating that the poly(A) tail is essential for PUM repressive activity. Thus, the mRNA with 3′ MALAT1 structure is resistant to PUM repression.

We examined whether an internal stretch of poly-adenosines could restore repression by PUMs to a similar degree as the 3′ poly(A) track. To do so, we introduced 20 or 60 adenosines immediately upstream of the MALAT1 triple helix to generate Nluc 3xPRE A20 MALAT and Nluc 3xPRE A60 MALAT reporters. These internal A20 and A60 tracts recruit PABPCs but are not substrates for deadenylases (15). The A20 track did not restore PUM repression and the A60 track supported a minor level of PUM repression in comparison to the poly-adenylated reporter **(Figure 1B**, compare 3xPRE A20 MALAT, 3xPRE A60 MALAT to Nluc 3xPRE). Therefore, robust repression required an exposed 3′ poly(A) tail, consistent with a PUM:CCR4-NOT deadenylation mechanism.

Given that poly(A) is necessary for PUM repression, we tested the potential role of the major cytoplasmic deadenylases. In addition to CCR4-NOT, the PAN deadenylase can catalyze poly(A) tail removal in human cells (14). The potential role of PAN in PUM repression has not been investigated previously. We therefore depleted the PAN2 and PAN3 subunits of the PAN deadenylase by RNA interference (RNAi) and tested the resulting effect on PUM repression of Nluc 3xPRE. Depletion of the PAN2 catalytic subunit was confirmed by western blot (**Figure 1C**); however, because an effective antibody to PAN3 is not available, we were unable to confirm its depletion. PAN2-PAN3 depletion did not reduce PUM repression; a modest increase was observed **(Figure 1D)**. In contrast, depletion of the CNOT1 subunit, which is the structural backbone of the CCR4-NOT complex, abolished PUM-mediated repression (37). We conclude that CCR4-NOT is necessary for PUM repression whereas PAN deadenylase is not.

### PUM1 associates with PABPC1 and PABPC4

The requirement of 3′ poly(A) tail for PUM repression suggested that poly(A)-binding proteins may participate in PUM repression. Previous work supports a connection of PUM and PABPC orthologs in budding yeast and *Drosophila* (45, 46). In addition, a high-throughput BioID analysis indicated that PUM1&2 can associate with PABPC1&C4 in cells (47). We therefore investigated the potential interaction of PUM1 with PABPCs using a co-immunoprecipitation assay. Endogenous PUM1 was immunoprecipitated from HCT116 cell extract, as confirmed by the enrichment for PUM1 in the immunoprecipitate, in comparison to the IgG negative control (**Figure 2A**). In contrast, the negative control GAPDH protein was not immunoprecipitated (**Figure 2A**). Both endogenous PABPC1 and PABPC4 coimmunoprecipitated with PUM1. Co-immunoprecipitation was performed in the presence of RNases A and One and loss of ribosomal RNAs confirmed RNA digestion (**Figure 2B**). We conclude that the PUM1 associates with PABPC1&C4 and is unlikely to be bridged by RNA. Minor signals for the translation initiation factors eIF4E and eIF4G were also detected in PUM1 immunoprecipitates (**Figure 2A**), suggesting that PUMs do not obstruct their interaction with PABPCs.

**Figure 2.**
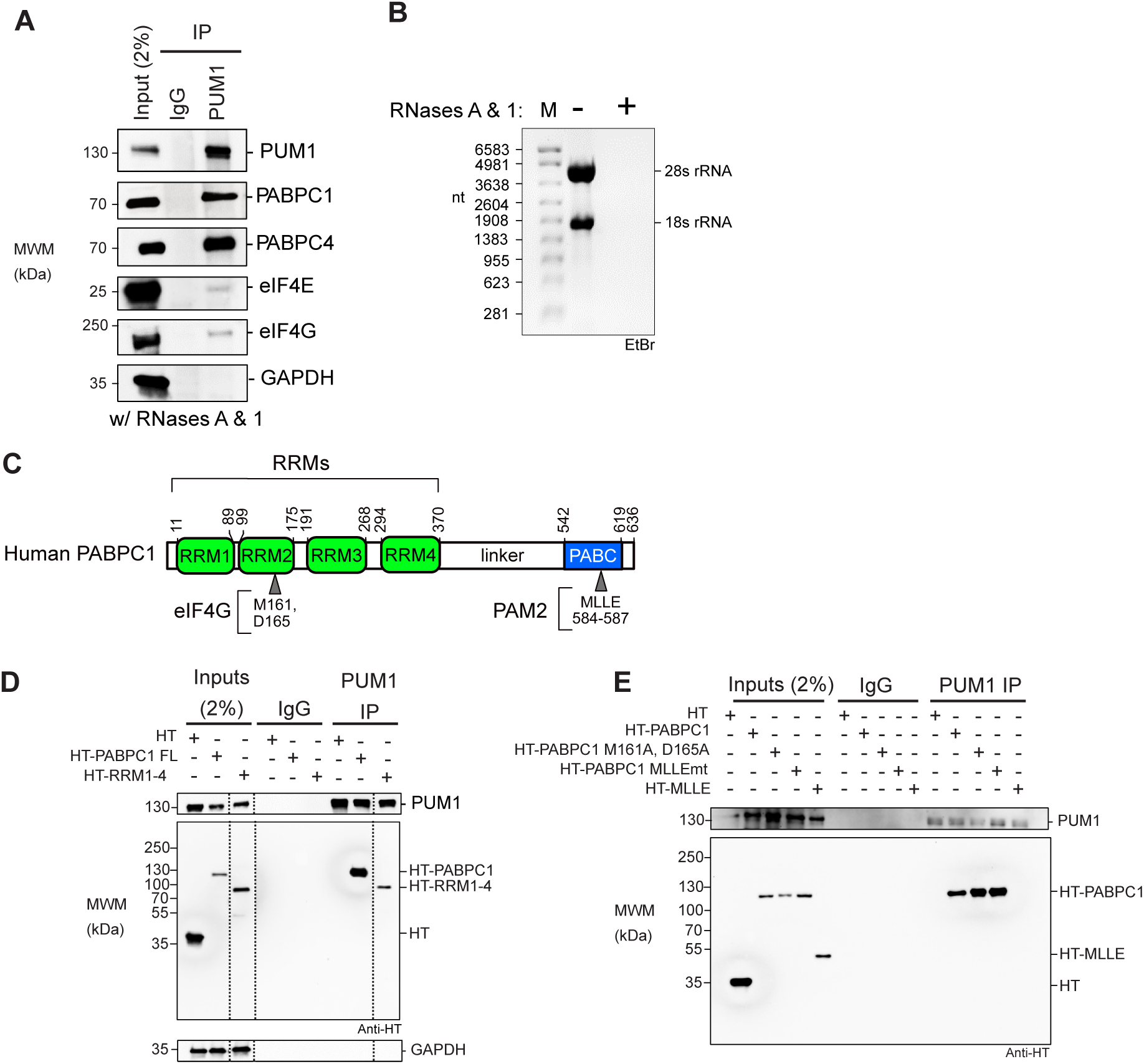
PUM1 interacts with endogenous PABPCs through the RRM domain independent of RNA. **A.** PABPC1 and PABPC4 co-immunoprecipitate with PUM1 from HCT116 cell extracts treated with RNases A and One. PUM1, eIF4E, eIF4G, PABPC1, PABPC4, and GAPDH were detected by western blot. **B.** Denaturing formaldehyde agarose gel analysis confirmed depletion of RNA in the HCT116 cell extracts before (-) or after treatment with RNase A and RNase One (+). The 18S and 28S ribosomal RNA bands, detected by ethidium bromide, are indicated on the right. **C.** Human PABPC1 domain architecture showing the N-terminal RRM domains (green) with the critical eIF4G binding site residues and the proline rich linker and C-terminal region PABC domain (blue) containing the MLLE motif residues important binding by PAM2-domain containing proteins. **D.** Western blot of the co-immunoprecipitation analysis of endogenous PUM1 with either HaloTag (HT), as a negative control, or HT-PABPC1 full-length (aa1-636) or HT-RRMs 1-4 (aa 1-370). All input samples were treated with RNase A and One. IgG beads and GAPDH served as negative controls. Dashed vertical lines in the panels indicate that the images were cropped to show relevant lanes. **E.** Western blot of PUM1 immunoprecipitates to detect association with either wild type full-length HT-PABPC1, or mutant versions wherein the eIF4G binding site is mutated (M161A and D165A) or the MLLE motif is mutated to (MLLEmt: M584G, L585A, L586A, and E587R), or the HT-MLLE domain (aa 542-636). All input samples were treated with RNase A and One.

To dissect the PUM1:PABPC1 association, we expressed PABPC1 constructs with N-terminal halotag (HT) fusions and assessed their interaction with endogenous PUM1. PABPC protein (**Figure 2C**) possesses four RNA recognition motifs (RRM1-4) that bind poly(A) (26, 48–52) and harbor the eIF4G binding site (**Figure 2C**). The RRM domain is joined by a proline-rich linker to the C-terminal PABC region containing the MLLE motif, which is recognized by multiple protein partners via their PAM2 motif (**Figure 2C**) (53). We found that both full-length HT-PABPC1 co-immunoprecipated with PUM1, whereas the negative control halotag or endogenous GAPDH did not (**Figure 2D**). The PABPC RRM1-4 fragment was sufficient for PUM1 association.

We then tested the effect of several PABPC mutations that prevent binding of its partners. Disrupting the eIF4G binding site (M161A, D165A) or the MLLE motif did not alter PUM1 association, indicating an interaction independent of these interfaces (**Figure 2E**)(54). These results show that PUM1 interacts in a distinct way compared to eIF4G or the known PABPC1 partners that recognize the MLLE motif (55). We also assessed the ability of PUM1 to associate with the PABC region containing the MLLE motif; no interaction was observed (**Figure 2E**).

From these results we conclude that PABPC1 RRMs are necessary and sufficient for interaction with PUM.

### PABPC1 and PABPC4 are necessary for PUM-mediated repression

The importance of the 3′ poly(A) tail for PUM repression and the physical association of PUM and PABPC prompted us to investigate PABPC as a potential PUM co-regulator. We first interrogated the predominant PABPC1 by performing RNAi knockdown in HCT116 cells (**Figure S2A**). PUM repression activity was measured using the Nluc 3xPRE reporter relative to the PRE mutant version. In this and subsequent experiments, we used a cotransfected internal control plasmid that expressed Fluc with a 3′ MALAT1 triple helix (Fluc MALAT) to minimize PABPC depletion effects on normalization. We observed that the knockdown of PABPC1 significantly reduced PUM repression compared to the negative control NTC siRNAs (**Figure S2B**). As an alternative strategy, we also used a cell line wherein the auxin inducible degron (AID) tag was appended to endogenous PABPC1, which yielded efficient and rapid depletion of PABPC1 when the auxin indole-3-acetic acid (IAA) was added to cells (30) (**Figure S2C,** compare with +IAA to without -IAA). Depletion of PABPC1-AID caused a statistically significant reduction in PUM repression compared to the untreated control (**Figure S2D**).

HCT116 cells express abundant PABPC1 and PABPC4 (**Figure 2A**) at a 3:1 ratio (37). We speculated that PABPC1 and PABPC4 may have overlapping or redundant functions in modulating PUM repression; therefore, we depleted PABPC1, PABPC4, or both simultaneously using AID and RNAi, respectively (**Figure 3A**) (30, 56, 57). Individually, depletion of PABPC1 or PAPBC4 caused a small reduction of PUM repression of the Nluc 3xPRE reporter (**Figure 3B**). Combined depletion of PABPC1 and PABPC4 eliminated repression (**Figure 3B**). Likewise, PABPC1&C4 depletion (**Figure 3C**) significantly reduced PUM repression of reporters controlled by the 3′ UTRs of natural PUM target mRNAs, FZD8 (**Figure 3D**) and CDKN1B (**Figure 3E**) (36). Depletion of PABPCs did not alter PUM1 and PUM2 levels. As expected, the non-polyadenylated internal control Fluc MALAT was unaffected by depletion of PABPCs (**Figure S2E**). Because PABPC1 is essential and others have reported decreased viability in response to extended PABPC-depletion in HeLa cells (29), we assessed cell health. With the short 24 hr depletion of PABPC1&C4, cell number and viability were unchanged (**Figure 3F & 3G**). Together, the results of this analysis demonstrate that PABPC1&C4 are necessary for PUM-mediated repression.

**Figure 3.**
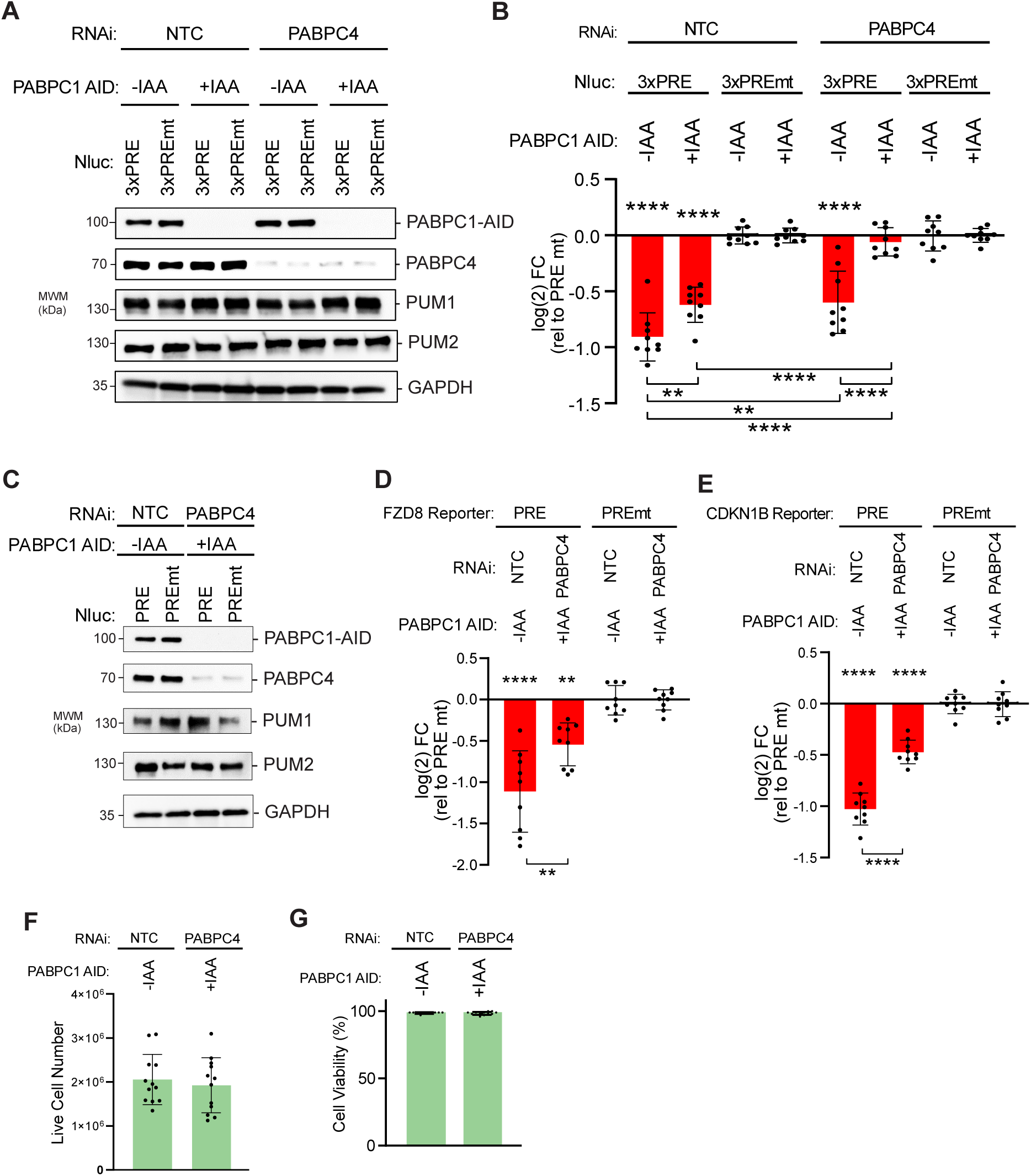
Poly(A)-binding proteins PABPC1 and PABPC4 are necessary for PUM-mediated repression. **A.** Western blot analysis confirms auxin induced depletion of degron-tagged PABPC1-AID and RNAi depletion of PABPC4 either individually or in combination in HCT116 cells. RNAi was performed by transfecting cells with either PABPC4 or non-targeting control NTC siRNAs. PABPC1-AID protein was depleted upon treatment with indole-3-acetic acid (+IAA), in comparison to vehicle only control (-IAA). GAPDH served as a loading control. **B.** PUM-mediated repression of Nluc 3xPRE reporter expression was measured in HCT116 cells, relative to the mutant PRE version. Individual and combined effects of PABPC4 RNAi and IAA induced depletion of PABPC1-AID on PUM repression were tested. n=9; 3 experiments, each with 3 biological replicates; +/- SD. For significance calling, p < 0.05 = *, p < 0.01 = **, p < 0.001 = ***, p < 0.0001 = **** based on ordinary one-way ANOVA and Tukey test for multiple comparisons. **C.** Western blot analysis of samples from a representative experimental replicate for experiments shown in Panels D and E. GAPDH served as a loading control and PUM1 and PUM2 protein levels were detected as controls. The effect of combined PABPC4 RNAi and PABPC1-AID depletion on PUM repression of Nluc reporters containing the 3′ UTRs of the natural, PRE-containing, PUM target mRNAs FZD8 in panel **D** and CDKN1B in panel **E** were measured, relative to matched PRE mutant versions. n=9; 3 experiments, each with 3 biological replicates; +/- SD. Live cell numbers (panel **F**) and viability (panel **G**) were measured in control or PABPC1-AID and PABPC4 RNAi depletion conditions from reporter assays shown in panel B. n=12; 3 experiments, each with 2 biological replicates, 2 technical replicates; +/- SD.

### PUM does not disrupt PABPC1 binding to mRNA

We considered the possibility that PUMs could cause poly(A) and PABPC dependent repression by displacing PABPC from the target mRNA. Such a mechanism has been proposed for other mRNA regulators (58). To test this hypothesis, we analyzed the interaction of PABPC1 with the PUM-repressed Nluc 3xPRE mRNA, in comparison to the unregulated mutant version, Nluc 3xPREmt. Endogenous PABPC1 was immunoprecipitated from cells expressing either reporter mRNA. After extensive washing, PABPC1 and associated mRNA were analyzed by western and northern blotting, respectively (**Figure 4A**). Both Nluc 3xPRE and 3xPREmt mRNAs were similarly enriched in the PABPC1 immunoprecipitates when normalized to inputs (**Figure 4B**). We conclude that PUM repression does not disrupt the interaction of PABPC1 with the target mRNA.

**Figure 4.**
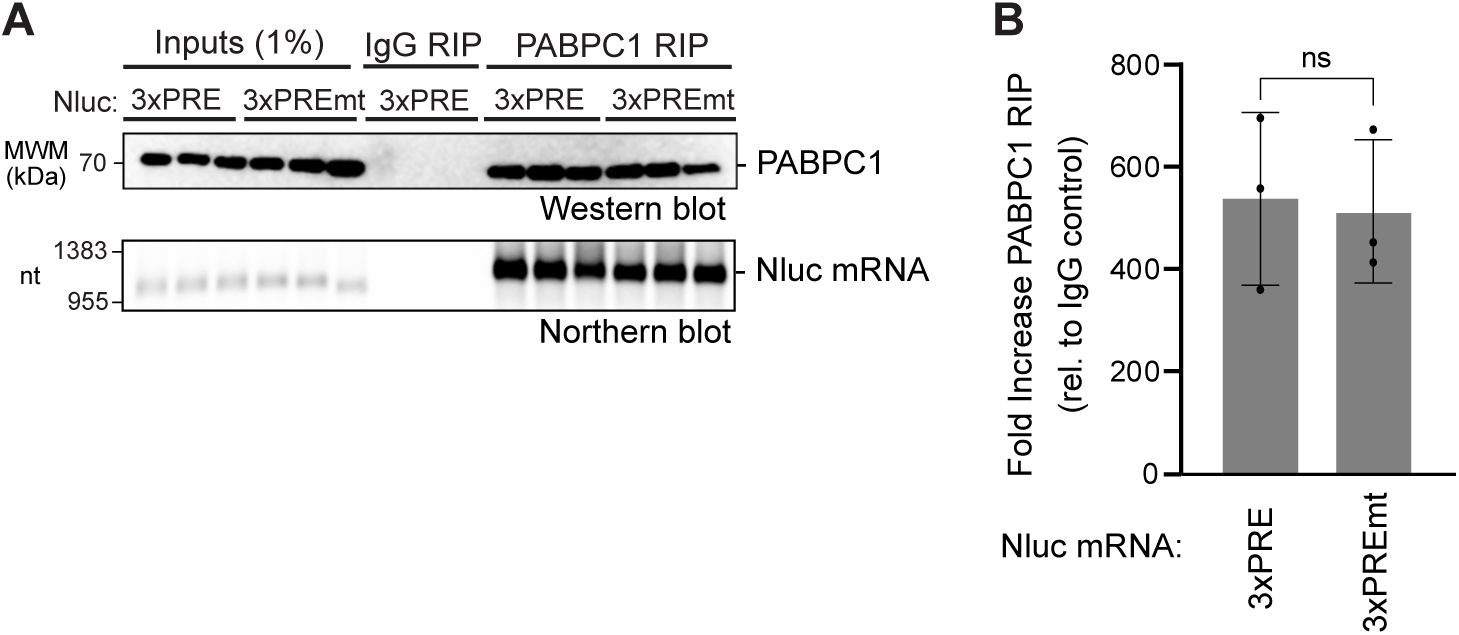
PUMs do not disrupt PABPC1 RNA binding. **A.** RNA co-immunoprecipitation (RIP) of endogenous PABPC1 with Nluc 3xPRE or 3xPREmt reporter RNAs was performed from HCT116 cell extracts. The top panel shows western blot detection of endogenous PABPC1 protein in the 1% input cell extract and 25% of the immunoprecipitate PABPC1 or negative control IgG RIP samples. The bottom panel shows detection of Nluc mRNA mRNAs by northern blotting. 1% of input and 75% of the RIP purified RNA samples were loaded on the denaturing formaldehyde-MOPS agarose gel, respectively. Three biological replicates were analyzed for each condition. **B.** Enrichment of Nluc 3xPRE or 3xPREmt reporter RNA in PABPC1 RIP samples was measured as fold increase in the mRNA relative to the IgG control. Nluc RNA level in each RIP sample was normalized to its level in the respective input sample. n=3 biological replicates; +/- SD. No significant (ns) difference in binding of each mRNA to PABPC1 was detected based on an unpaired student’s t-test.

### PABPC and PUM have opposing effects on the rate of mRNA degradation

PUM1&2 repress target mRNAs by accelerating their degradation (35–37, 59). We hypothesized that PABPC-driven stabilization likely underlies its role in PUM repression. To test how PABPC affects PUM-mediated mRNA degradation, we measured the effect of PABPC1&C4 depletion on the stability of PRE-containing reporter mRNA in comparison to the non-target version with mutated PREs. The optimized experimental strategy and timeline is shown in **Figure 5A**. This experiment used a Nluc reporter with six PREs, providing enhanced response to PUM repression, which is alleviated by depletion of PABPC1&C4 (**Figure S3A**). Using the Tet-Off promoter, transcription of the reporter was inhibited in the presence of doxycycline and activated by its removal to produce a pulse of nascent transcript. Re-addition of doxycycline shut off transcription, and RNA samples were collected over an 8 hour time course (**Figure 5A**). The RNA was analyzed by northern blot and the half-life of the mRNAs was calculated, using data collected in three independent experiments (see **Figure 5B** for representative experiment and **Figure S3** for additional replicates). To ensure efficient depletion of PABPC1, we engineered a new auxin inducible degron system in the HCT116 cell line wherein PABPC1 has an AID degron fused to its C-terminus (**Figure S3B and S3C**). To mediate specific degradation of the fusion, the auxin signaling F-box protein from *Arabidopsis thaliana*, AFB2, was inserted into the AAVS1 “safe harbor” locus (60).

**Figure 5.**
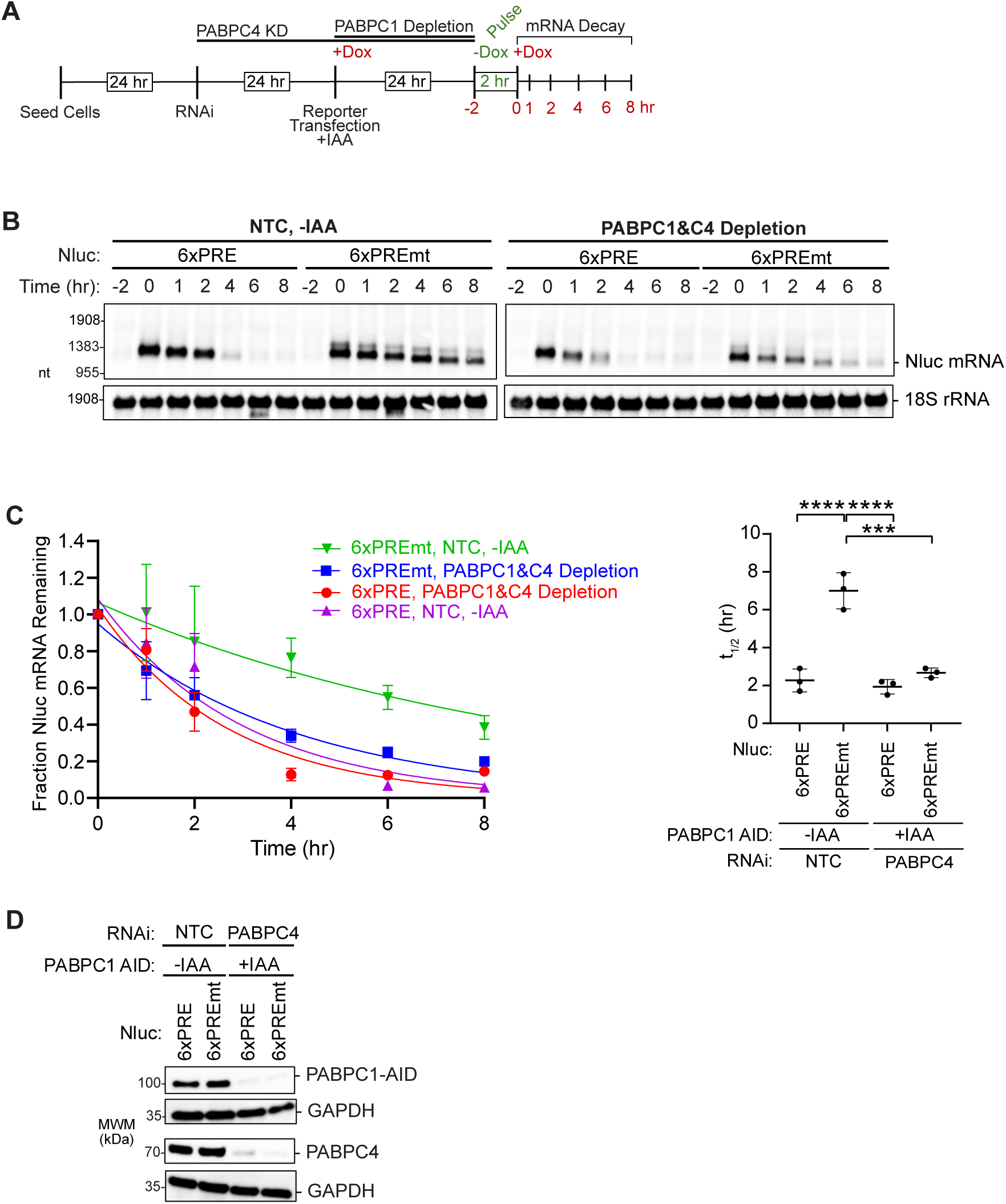
PABPC1&4 depletion accelerates mRNA degradation independent of PUM activity. **A.** Experimental strategy for measuring mRNA decay rates in response to PUM repression and depletion of PABPC1-AID and PABPC4. After RNAi knockdown of PABPC4 and auxin (+IAA) induced degradation of PABPC1-AID, the tet-off regulated expression of Nluc 6xPRE or Nluc 6xPREmt produced a pulse of nascent Nluc mRNA in HCT116 cells. Time course shows the procedure for RNAi induced PABPC4 depletion and auxin (+IAA) induced PABPC1-AID depletion. The timeline for transfection of the Tet-off Nluc reporters is shown, along with the suppression of the reporters by doxycycline (+Dox) and transcriptional pulse caused by its removal (-Dox) for 2 hours. RNA samples were collected before the pulse (-2hrs) and at 0, 1, 2, 4, 6, and 8 hours post-pulse to measure decay rates by northern blot. **B.** Tet-off transcription shut-off was performed to compare the half-lives of the Nluc 6xPRE and 6xPREmt reporter mRNAs under NTC or PABPC1&C4 depletion. A representative northern blot of Nluc reporters and the 18S ribosomal rRNA internal control is shown. Replicate blots are shown in Supplemental Figure S3. **C.** Decay rates of Nluc 6xPRE and Nluc 6xPREmt in response to depletion of PABPC1 and PABPC4 (left). The fraction of each Nluc mRNA remaining, normalized to internal control 18S rRNA, is plotted relative to time (hours) after inhibition of transcription. First order exponential decay trend lines, calculated by non-linear regression analysis, are plotted for each experimental condition from three experimental replicates. n=3; +/- SD. On the right, mean Nluc mRNA half-lives from the experimental replicates are compared. n=3; +/-SD. For significance calling, p < 0.05 = *, p < 0.01 = **, p < 0.001 = ***, p < 0.0001 = **** based on ordinary one-way ANOVA and Tukey test for multiple comparisons. **D.** A representative western blot confirming depletion of PABPC1 and PABPC4 by AID and RNAi, respectively.

As expected for a PUM target, the Nluc 6xPRE mRNA decayed with a significantly shorter half-life (3-fold) than the mutant version in the control condition (**Figure 5B and 5C**, compare 6xPRE vs 6xPREmt in the NTC, -IAA condition). Depletion of PABPC1&C4, verified in **Figures 5D and S3E**, abrogated the acceleration of mRNA decay by wild type PREs (**Figure 5B and 5C**, compare 6xPRE vs 6xPREmt in the PABPC4 RNAi, +IAA condition). This loss of PUM-mediated degradation was due to the significant destabilization of the normally stable Nluc 6xPREmt mRNA (**Figure 5C**, 2.6-fold decrease in 6xPREmt mRNA half-life). Whereas the half-life of the Nluc 6xPRE mRNA was rendered indistinguishable between the control and PABPC depletion conditions (**Figure 5C**). Taken together, these results show that the principal effect of PABPCs is mRNA stabilization. Without PABPC1&C4, PRE-dependent acceleration contributes little because basal stability is reduced. We conclude that PUMs and PABPCs have opposing effects on mRNA decay. PABPCs stabilize mRNAs, both regulated and unregulated by PUMs, whereas PUMs cause degradation of PRE-containing mRNAs. We conclude that PABPCs are necessary for PUM repression because they stabilize mRNAs.

### PUMs and PABPCs have opposing effects on the levels of natural mRNAs

To corroborate our observations on the opposing effects of PUMs and PABPCs, we analyzed several natural PUM target mRNAs including ITGA2 and SMPDL3A, both of which have two PREs in their 3′ UTR and are bound and repressed by PUM1 and PUM2 (36, 37). Indeed, depletion of PUM1&2 increased the levels of ITGA2 and SMPLD3A mRNAs relative to non-targeting control siRNAs, as measured by RTqPCR (**Figure 6A**). PUM depletion had no effect on the non-target GAPDH mRNA or the noncoding MALAT1 RNA (**Figure 6A**). In contrast, depletion of PABPC1&C4 reduced the levels of PUM target mRNAs ITGA2 and SMPDL3A (**Figure 6B**). Similarly, the non-target GAPDH mRNA was reduced in response to PABPC depletion. The non-adenylated MALAT1 ncRNA was not affected. Depletion of PUMs and PABPCs in these experiments were confirmed by western blot (**Figure 6C, 6D**). Overall these results support the conclusion that PUMs and PABPCs have opposing effects on levels of endogenous natural mRNAs. PUMs cause degradation of PRE-containing mRNAs, whereas PABPCs stabilize poly(A) mRNAs.

**Figure 6.**
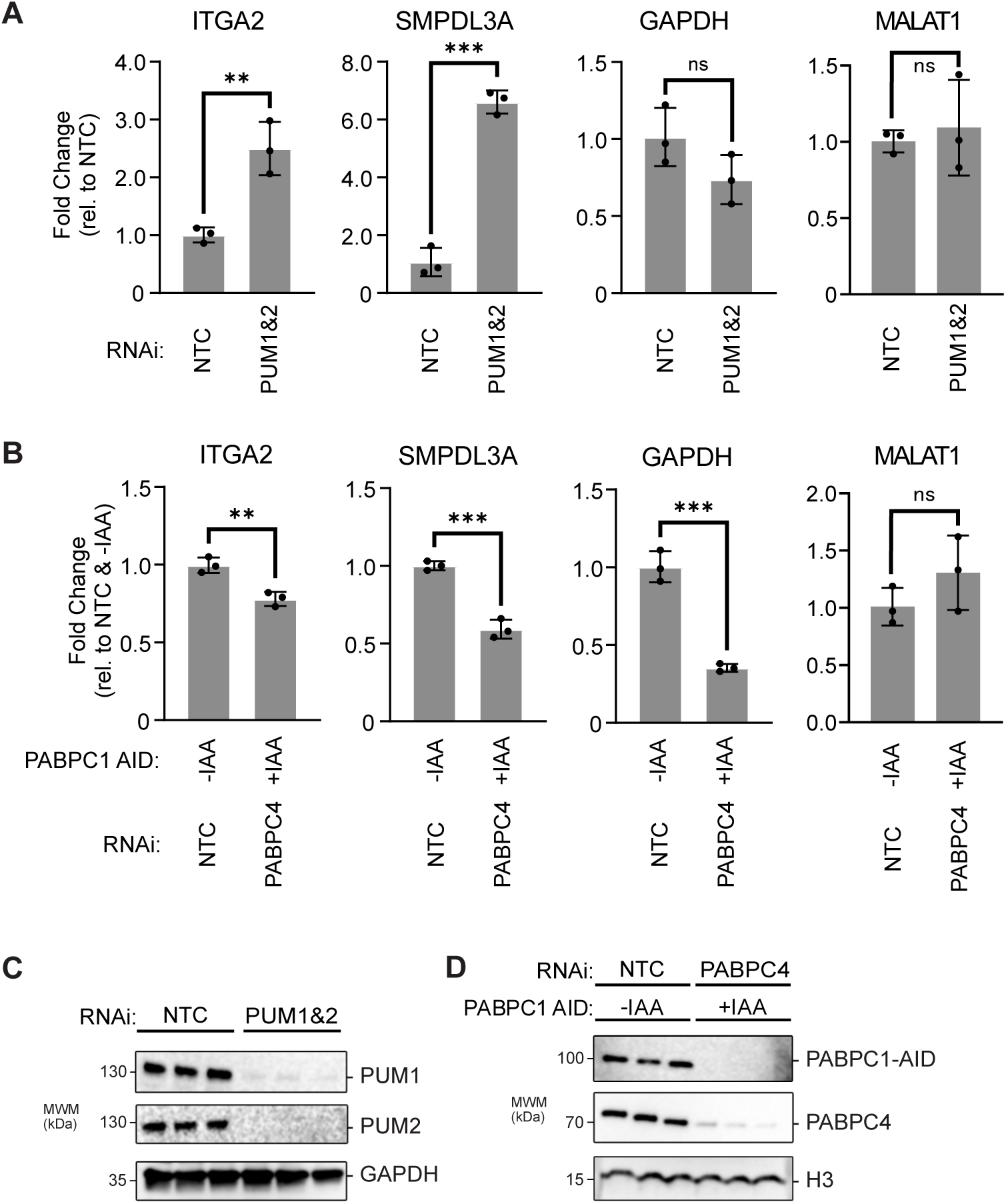
PUMs and PABPCs have opposing effects on mRNA stability. **A.** Expression levels of the PRE-containing PUM target mRNAs ITGA2 and SMPDL3A were measured by RTqPCR in RNA samples purified from HCT116 cells wherein PUM1 and PUM2 were depleted by RNAi. Changes in mRNA levels were determined relative to cells treated with the non-targeting control siRNAs (NTC). The non-targeted GAPDH mRNA and the non-adenylated MALAT1 non-coding RNA served as controls. **B.** Expression levels of the PRE-containing PUM target mRNAs ITGA2 and SMPDL3A were measured by RTqPCR in RNA samples purified from HCT116 cells wherein PABPC1-AID and PABPC4 were depleted by auxin treatment (+IAA) and RNAi, respectively. Changes in mRNA levels were determined relative to cells treated with vehicle only (-IAA) and the NTC siRNA. The non-targeted GAPDH mRNA and the non-adenylated MALAT1 non-coding RNA served as controls. In both panels A and B, transcript level was measured by RTqPCR, normalized to 18S rRNA, and plotted as fold change relative to the non-depleted control condition. n=3 biological replicates, ± SD. For significance calling, p < 0.05 = *, p < 0.01 = **, p < 0.001 = ***, p < 0.0001 = ****, ns = not significant based on unpaired two-tailed t-tests. **C.** Western blot analysis confirmed RNAi depletion of PUM1 and PUM2 in biological replicates in panel A. GAPDH served as a loading control. **D.** Western blot analysis of PABPC1-AID and PABPC4 confirmed their depletion by auxin induced degradation and RNAi, respectively, in samples from panel B. Histone H3 served as a loading control.

### Over-expression of PABPC1 opposes PUM-mediated repression

To further interrogate the model that PUMs and PABPCs have opposing effects on mRNA regulation, we tested the effect of increasing PABPC levels on PUM repression. To do so, increasing amounts of an HT-PABPC1 expression plasmid were transfected into HCT116 cells along with the Nluc 3xPRE or Nluc 3xPREmt reporters and the Fluc MALAT control plasmid.

Western blotting confirmed HT-PABPC1 expression and we measured up to 3-fold excess over endogenous PABPC1 levels (**Figure 7A**). Exogenous HT-PABPC1 inhibited PUM repression of the Nluc 3xPRE in a dose-dependent manner (**Figure 7B**). We confirmed PUM1 and PUM2 protein expression levels by western blot, which were unaffected by PABPC1 over-expression (**Figure 7A**). As expected, the activity from the Fluc MALAT internal control was unaffected by PABPC1 over-expression (**Figure S2F**). These results provide additional evidence that PABPC1 can oppose PUM-mediated repression. They also indicate that the abundance of PABPC1 in these cells is not saturated.

**Figure 7.**
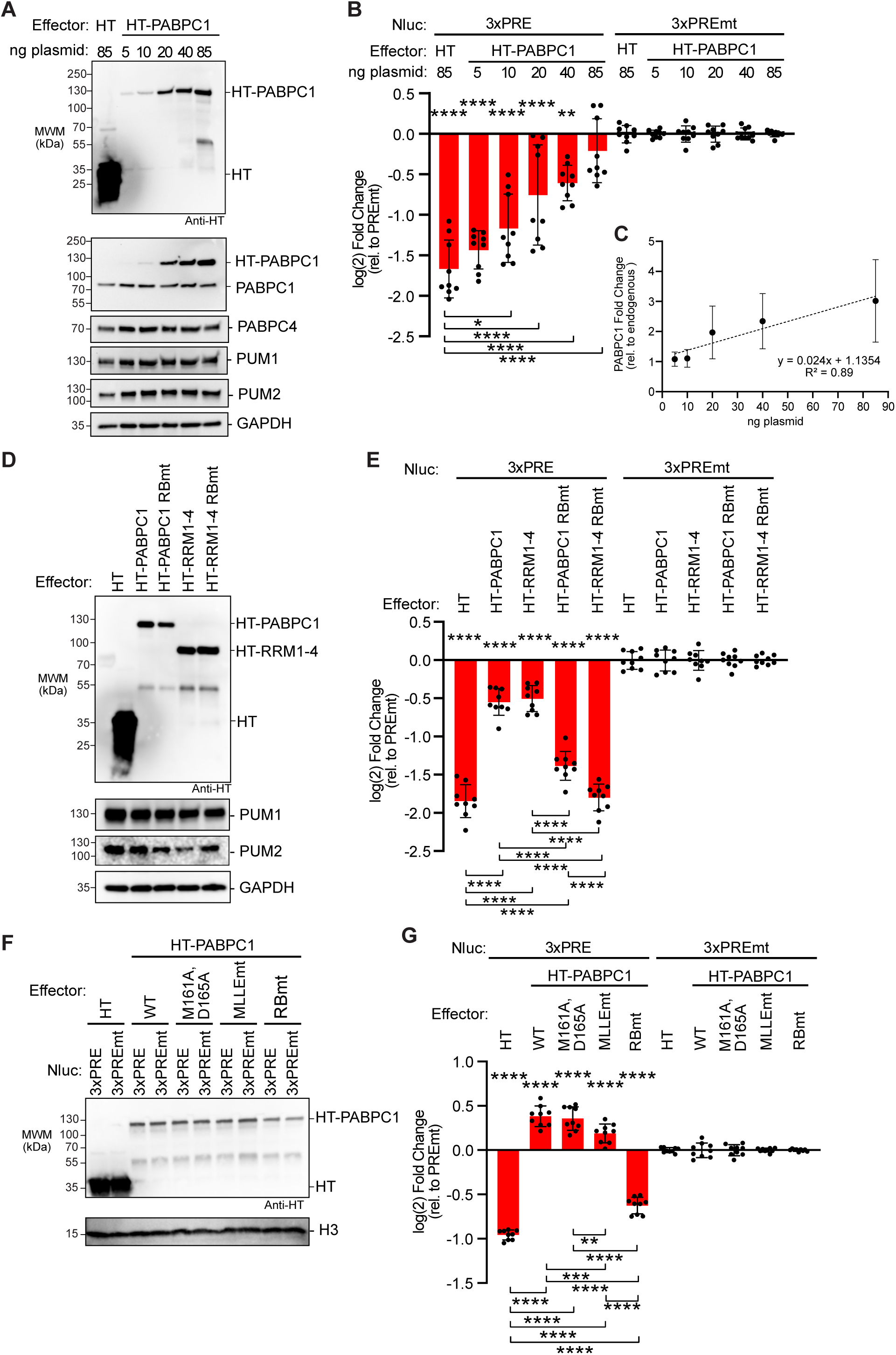
PABPC over-expression alleviates PUM-repression and requires RNA binding. **A.** Western blot analysis of halotag (HT) and HT-PABPC1 titration for samples from a representative experimental replicate of panel B. The amount of transfected plasmid for each effector is shown at the top. GAPDH served as a loading control. PUM1 and PUM2 were detected as an additional control. **B.** Reporter assay showing the effect of HT-PABPC1 over-expression on PUM repression of the Nluc 3xPRE reporter in wild type HCT116 cells, calculated relative to the Nluc 3xPREmt reporter. Halotag served as a negative control. n=9; 3 experiments, each with 3 biological replicates; +/- SD. For significance calling, p < 0.05 = *, p < 0.01 = **, p < 0.001 = ***, p < 0.0001 = **** based on ordinary one-way ANOVA and Tukey test for multiple comparisons. Significance indicated above the X-axis indicates relative to 3xPREmt reporter, whereas significance calling shown below is indicated by the respective brackets. **C.** Graph of the fold change in HT-PABPC1 exogenous expression over endogenous PABPC1 levels calculated from three experimental replicates including the western blot shown in panel A. n=3; +/- SD. **D.** Western blot analysis of halotag (HT), HT-PABPC1 full-length, HT-PABPC1 full-length RNA-binding mutant (RBmt), HT-RRM1-4 domains, and HT-RRM1-4 RBmt samples taken from a representative experimental replicate of panel E. GAPDH served as a loading control. PUM1 and PUM2 were detected as controls. **E.** Reporter assay showing effect of HT-PABPC1 full-length, HT-RRM1-4 domains, and RNA-binding mutants versions when over-expressed on PUM repression of the Nluc 3xPRE reporter in HCT116 cells. n=9; 3 experiments, each with 3 biological replicates; +/- SD. Halotag served as a negative control. **F.** Western blot analysis of over-expressed HT-PABPC1 on PUM repression and the effect of the mutation of the eIF4G binding site mutant (M161A, D165A), or the MLLE motif (MLLEmt), or the RNA-binding defective mutant (RBmt) from a representative experimental replicate of panel G. H3 served as a loading control. PUM1 and PUM2 were detected as additional controls. **G.** Reporter assay showing effect of HT-PABPC1 full-length mutants on PUM repression in wild type HCT116 cells. n=9; 3 experiments, each with 3 biological replicates; +/- SD. Halotag served as a negative control.

Because the over-expressed PABPC1 had an N-terminal halotag moiety, we considered the possibility that this might alter PABPC1 activity. Therefore, we performed an identical experiment using untagged PABPC1. The results confirm that increasing PABPC1 level inhibits PUM-mediated repression **(Figure S4A and S4B)**; therefore, the fusion to halotag did not alter PABPC1 function.

### Binding of PABPC1 to RNA is essential for its ability to oppose PUM repression

We utilized the over-expression assay to determine which features of PABPC1 are necessary and sufficient to inhibit PUM. We found that the four RRMs were capable of inhibiting PUM repression to the same degree as full length PABPC1 (**Figure7D, 7E,** compare HT-PABPC1 to HT-RRM1-4), suggesting that the RNA-binding function of PABPC1 was important. To assess the role of PABPC RNA-binding activity, we tested an RNA-binding-defective PABPC1 (RBmt; RNP1 mutations in RRM1-4), which failed to inhibit PUM repression, demonstrating the requirement for RNA binding. These mutations were previously shown to abolish RNA-binding by *Saccharomyces cerevisiae* ortholog PAB1(48, 62–63). We found that the mutations inactivated the ability of full length PABPC1 or RRM1-4 to inhibit PUM repression (**Figure 7D, 7E**). These results demonstrate that the RNA-binding domain and the ability of PABPC1 to bind poly(A) RNA are essential for opposing PUM repression.

We analyzed the role of known PABPC1 protein-protein interactions by mutating the eIF4G binding site (M161A, D165A) or the MLLE motif (MLLEmt) **(Figure 2C)** (15). When over-expressed, these PABPC1 mutants inhibited PUM repression to a similar extent as wild type PABPC1, whereas the RNA-binding mutant PABPC1 was inactive (**Figure 7F, 7G**). Thus, binding of PABPC1 to its protein partners is not necessary to inhibit PUM.

### Multiple PABPC paralogs can oppose PUM repression

To determine if PABPC1 is unique in its ability to inhibit PUM repression, we tested several PABPC paralogs, including PABPC3, PABPC4, and PABPC5 (**Figure S4C**). PABPC3 and PABPC4 share 92% and 75% identify with PABPC1, respectively. PABPC5 shares four RRMs with 65% identity to PABPC1, yet lacks the C-terminal linker and PABC domain containing the MLLE motif. Each PABPC paralog inhibited PUM repression when over-expressed (**Figure S4D-G**). Even the most divergent PABPC5 inhibited PUM activity; underscoring the importance of the PABPC RRMs in this activity. Thus, multiple PABPCs are capable of opposing PUM repression activity.

### Increased PABPC1 does not prevent binding of PUM to target mRNA

We considered the possibility that over-expressed PABPC may prevent PUM repression by interfering with binding of PUM to its target mRNA. We examined this by immunoprecipitating PUM1 and detecting associated Nluc 3xPRE mRNA by northern blot (i.e. RNA coimmunoprecipitation, RIP, assay) in the presence of over-expressed HT-PABPC1 **(Figure 8A)**. Over-expression of HT-PABPC1 stabilized the Nluc 3xPRE mRNA, increasing its abundance by 5-fold relative to cells expressing HT alone (**Figure 8B**, Inputs). This increase carried over in the PUM1 immunoprecipitates, wherein more Nluc 3xPRE RNA was bound to PUM1 relative to the HT negative control (**Figure 8A**). Fold enrichment of the mRNA was calculated relative to the IgG negative control RIP samples. When normalized to the corresponding input levels of Nluc 3xPRE, we did not detect a statistically significant difference in PUM1-associated Nluc 3xPRE mRNA in the presence of overexpressed PABPC1 (**Figure 8C**). Based on these results, we conclude that over-expressed PABPC1 does not disrupt the interaction of PUM1 with its target mRNA.

**Figure 8.**
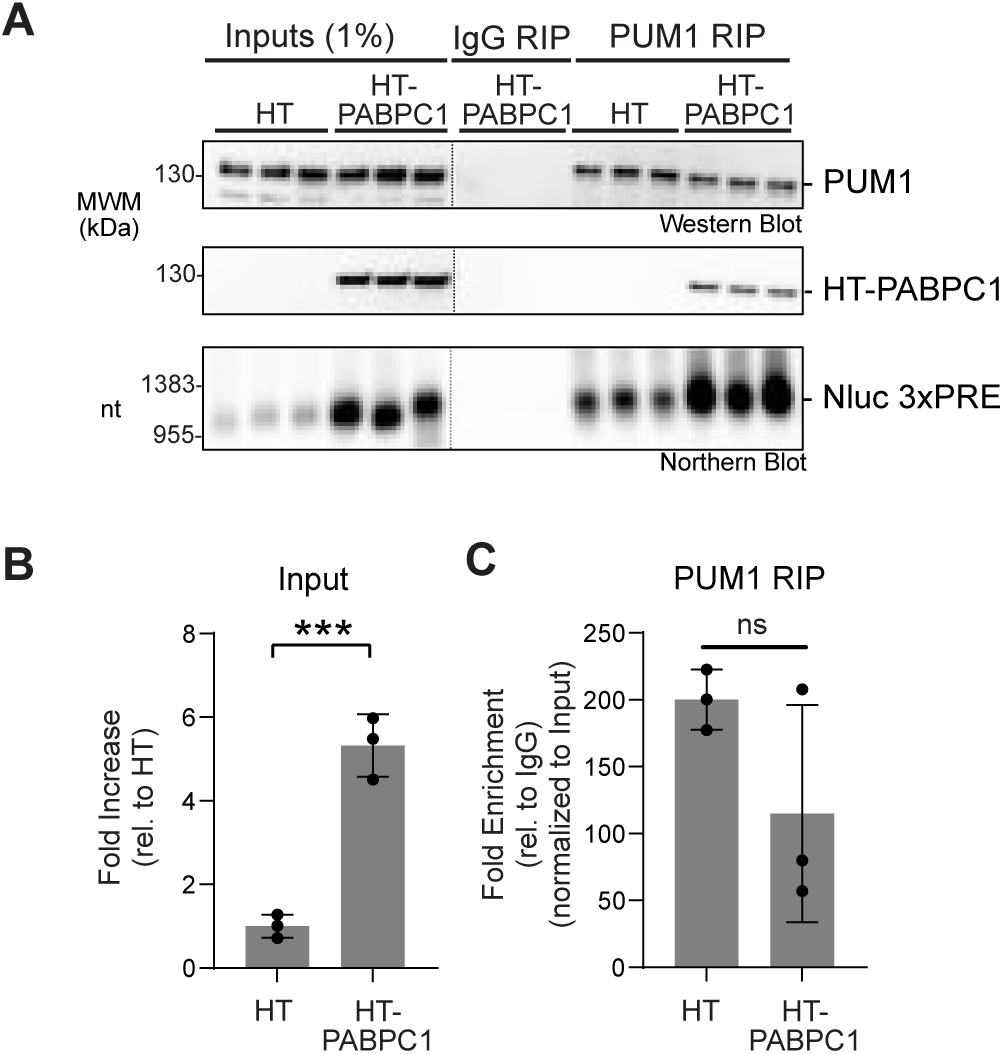
PABPC1 does not disrupt PUM binding to Nluc 3xPRE mRNA. **A.** Co-immunoprecipitation analysis of PUM1 binding to the Nluc 3xPRE reporter in the presence of over-expressed halotag (HT) or HT-PABPC1. The top two panels show western blot detection of PUM1 protein and HT-PABPC1 in the 2% of the input HCT116 cell extracts the and 25% of the PUM1 RNA co-immunoprecipitates (RIP) from three biological replicate samples per condition. The bottom panel shows detection of Nluc mRNA by northern blotting. For northern blots, 1% of input samples and 75% of the RIP samples were loaded on the formaldehyde-MOPS agarose gel, respectively. Co-immunoprecipitation with IgG beads served as a negative control. **B.** Fold increase of the Nluc 3xPRE mRNA in the input samples from cells expressing HT- PABPC1 are plotted relative to the HT control, based on data in panel A. n=3 biological replicates, +/- SD. For significance calling, p < 0.001 = *** based on an unpaired student’s t-test. **C.** Fold enrichment of the Nluc 3xPRE mRNA in PUM1 RIP samples was measured relative to the IgG control. Importantly, the mRNA levels in each RIP sample was normalized to that present in the respective input samples. n=3 biological replicates, +/- SD. For significance calling, the difference between HT and HT-PABPC1 was not significant (p > 0.05 = ns) based on an unpaired student’s t-test.

### Increased PABPC1 stabilizes target mRNAs

Based on the observation that PABPC1 increased the level of Nluc 3xPRE in Figure 8, we next examined the effect of increased PABPC1 on degradation of both PUM target mRNA and non-target mRNAs. To do so, we measured Nluc 6xPRE and 6xPREmt mRNA levels in cells that over-expressed HT-PABPC1 (**Figure 9A**). Expression of HT-PABPC1 relative to endogenous PABPC1 was confirmed by western blot (**Figure 9B**). Quantitation of the mRNA levels in each condition revealed that over-expressed PABPC1 significantly increased Nluc 6xPRE mRNA levels, relative to over-expressed HT control protein, and to a lesser degree also increased Nluc 6xPREmt mRNA (**Figure 9C**). In effect, over-expressed PABPC1 alleviated the ability of PUM to reduce the Nluc 6xPRE mRNA levels relative to the mutant reporter (**Figure 9D**) consistent with the effect on PUM repression of Nluc protein expression in Figure 7. Poly(A) tails appeared longer with PABPC1 over-expression, consistent with PABPC1 antagonizing deadenylation (**Figure 9A**).

**Figure 9.**
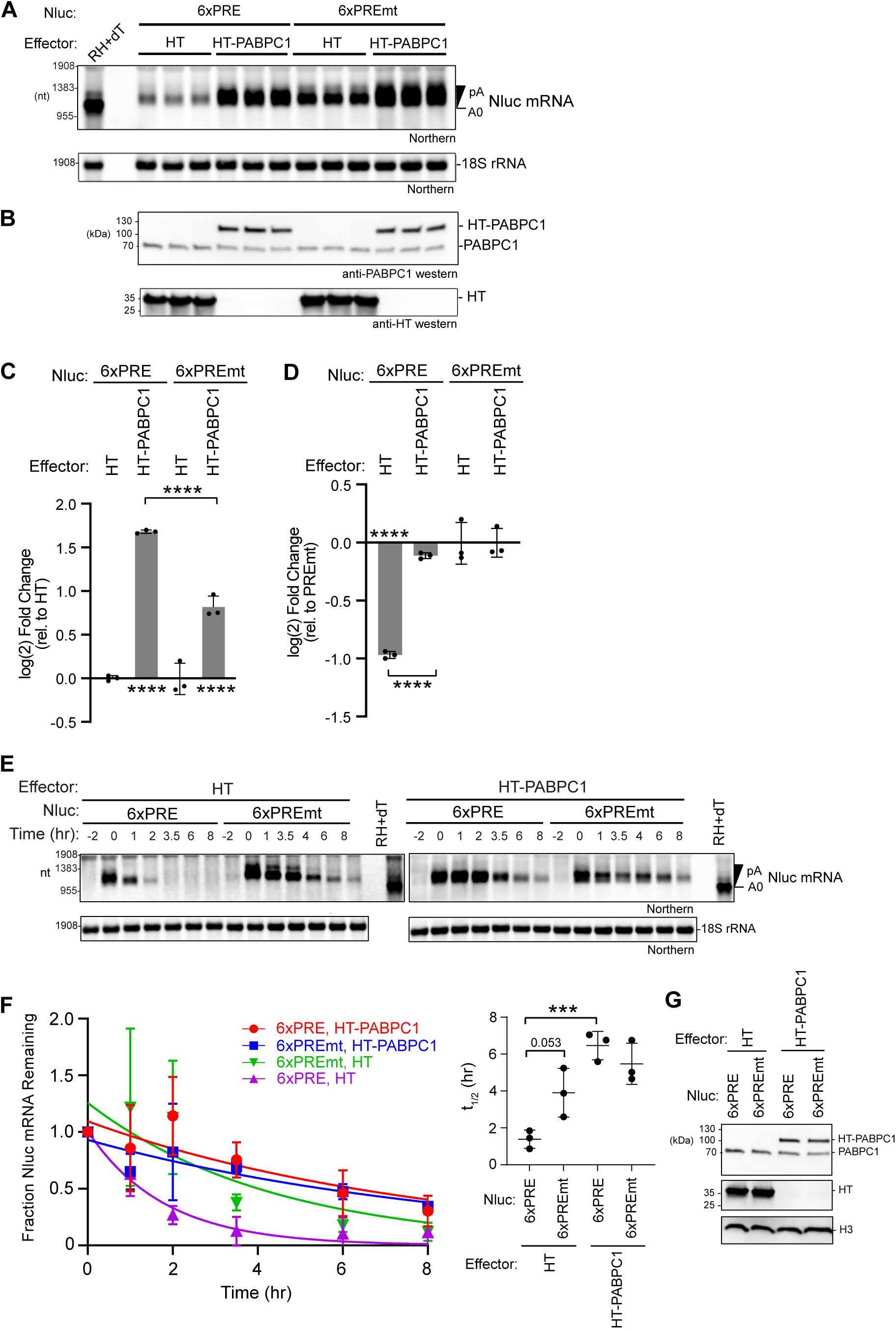
PABPC1 over-expression stabilizes mRNAs, blocking PUM-mediated mRNA degradation. **A.** Northern blot analysis of Nluc reporter mRNA containing either six wild type PREs (6xPRE) or mutant PREs (6xPREmt) in HCT116 cells transfected with halotag (HT) or HT-PABPC1 expression plasmids. 18S rRNA served as a loading control, and ethidium bromide staining confirmed RNA integrity and loading. Three biological replicate samples are shown for each condition. As a marker for the Nluc mRNA with the poly(A) tail removed (A0), one RNA sample from the HT control was treated with oligo-dT15 and RNase H (RH+dT). Poly-adenylated (pA) and deadenylated (A0) Nluc species are indicated on the right. **B.** Western blot analysis of effector protein expression. PABPC1 antibody detected both endogenous PABPC1 and over-expressed HT-PABPC1, while HT antibody confirmed the expression of HT. **C.** Fold changes of either Nluc 6xPRE or Nluc 6xPREmt mRNA levels from panel A in response to over-expressed HT-PABPC1 were calculated relative to their respective HT negative controls. Data represent mean values ± SD, n = 3 biological replicates. **D.** Fold changes of either Nluc 6xPRE mRNA levels from panel A in response to over-expressed HT-PABPC1 or the negative control HT were calculated relative to their respective or Nluc 6xPREmt negative controls. Data represent mean values ± SD, n = 3 biological replicates. For significance calling, p < 0.05 = *, p < 0.01 = **, based on ordinary one-way ANOVA and Tukey test for multiple comparisons. **E.** Tet-off transcription shut-off was performed to compare the half-lives of the Nluc 6xPRE and 6xPREmt reporter mRNAs in response to over-expressed HT-PABPC1 or negative control halotag (HT). A representative northern blot of Nluc reporters and the 18S ribosomal rRNA internal control is shown. Replicate blots are shown in Supplemental Figure S5. **F.** Decay rates of Nluc 6xPRE and Nluc 6xPREmt in response to over-expression of HT- PABPC1, in comparison to HT. The fraction of each Nluc mRNA remaining, normalized to 18S rRNA, is plotted relative to time (hours) after inhibition of transcription. First order exponential decay trend lines, calculated by non-linear regression analysis, are plotted for each experimental condition from three experimental replicates. n=3; +/- SD. On the right, mean Nluc mRNA half-lives from the experimental replicates are compared. n=3; +/-SD. For significance calling, p < 0.05 = *, p < 0.01 = **, p < 0.001 = ***, based on ordinary one-way ANOVA and Tukey test for multiple comparisons. **G.** A representative western blot confirming expression of HT-PABPC1 relative to endogenous PABPC1 using anti-PABPC1 antibody in each condition used for mRNA decay analysis. Western blot also confirmed expression of the HT protein. Histone H3 served as a loading control.

We then measured the effect of increasing PABPC1 on the rate of decay of reporter mRNAs using the Tet-Off regulated transcriptional pulse approach. Over-expressed HT-PABPC1 stabilized both the Nluc 6xPRE reporter mRNA relative to the negative control HT (**Figure 9E-9G,** with data from experimental replicates shown in **Figure S5**). As a result, PUM-mediated degradation of the PRE reporter was alleviated. These results show that increased PABPC1 preferentially blocks the ability of PUMs to effectively initiate mRNA decay.

### PABPC levels vary across normal human tissues

Our experimental results demonstrate that the abundance of PABPC protein can tune the responsiveness of mRNAs to RBP-directed regulation. We assessed whether such variation may naturally occur in physiological contexts by analyzing PABPC RNA and protein levels in different human tissues. First, we interrogated RNA-seq data in the GTEx database from normal human tissues (version 10, Gencode version 39, including 54 tissues from 946 donors and 19788 samples). The data demonstrate that median PABPC1 mRNA levels vary by 17-fold across tissues, with maximal expression in esophagus, vagina, ovary, and salivary gland and minimal expression in kidney, skeletal muscle, heart, and brain (**Figure S6A**). Similarly, median PABPC4 mRNA levels vary by 15-fold with maximal expression in skeletal muscle, pancreas, ovary and testes, and minimal expression in liver, kidney, blood, and brain (**Figure S6B**).

To further examine PABPC1 expression patterns, we performed western blot of available normal human tissues. It is important to recognize that these western blots are derived from individual specimens, whereas the GTEX mRNA data is derived from hundreds of donors. PABPC1 was detected in the testis, ovary, kidney, and brain, but not other tissues (**Figure S7A**). The signal for this PABPC1 antibody was low and crossreactivity with other proteins was observed. Therefore, we obtained another antibody that recognizes the PABPC1, PABPC3 and PABPC4 paralogs, which are indistinguishable based on their near identical 70-72 kDa molecular weights. These PABPCs were detected in the testis, ovary, skeletal muscle, kidney, and brain, but not the other tissues (**Figure S7B**). In the same manner, we analyzed PABPC4 protein levels using a specific antibody (**Figure S7C**). The highest levels of PABPC4 were detected in testis, followed by ovary, skeletal muscle, and intermediate levels in kidney, lung, and brain. PABPC4 was not detected in the liver, heart, small intestine, or spleen.

From these data, we conclude that the expression levels of PAPBC1 and PABPC4 vary widely across human tissues. PABPC protein levels are likely controlled by a combination of transcriptional, post-transcriptional, and post-translational mechanisms that remain to be determined. The implications of these differences, coupled with our experimental analysis, provide support for the hypothesis that PABPC abundance may naturally modulate mRNA regulation by cis- and trans-acting regulatory mechanisms.

## DISCUSSION

The results of this study reveal the central role that the 3′ poly(A) tail and PABPC proteins play in modulating the stability of mRNAs and their response to regulation. We found that the poly(A) tail is essential for PUM repression and must be at the 3′ end of the target mRNA, where it is accessible to shortening by the CCR4-NOT deadenylase complex. Our results reaffirm the essential role of CCR4-NOT in the mechanism of PUM-mediated repression (35, 37). In contrast, the PAN deadenylase does not appear to be required. This result suggests that the coordinated action PAN and CCR4-NOT may be unnecessary for PUM to promote degradation of target mRNAs.

Replacing the poly(A) tail of a PRE-containing target mRNA with the MALAT1 triple helix RNA structure renders the mRNA resistant to PUM:CCR4-NOT mediated repression. Indeed, the MALAT1 structure was reported to support translation and resist degradation by 3′ exoribonucleases (42–44). We also observed that inserting internal poly(A) tracts sufficient for binding PABPC molecules did not restore repression in the context of the PRE-containing mRNA with a MALAT1 structure at the 3′ end. Thus, PABPC association with the target mRNA is, in itself, not adequate to support PUM repression. Instead, the protection of poly(A) by PABPC is predicated on its 3′ end positioning and accessibility to deadenylation. The observed requirement of an accessible 3′ poly(A) tail is consistent with previous surveys of natural PUM target RNAs, which identified hundreds of PRE-containing, poly-adenylated mRNAs and ncRNAs that are degraded by PUMs (36, 37).

Our results reveal that PABPC is necessary for PUM repression. Cultured cells typically express both PABPC1 and PABPC4, which appear to act redundantly in this context (29, 30). Both PABPCs must be depleted to alleviate PUM repression. The results indicate that PABPCs and PUMs have opposing effects on mRNA stability. Under normal conditions, PABPCs generally stabilize mRNAs. For PRE-containing mRNAs, PUM must overcome the stabilizing activity of PABPCs to accelerate degradation. Experimental depletion of PABPCs renders the poly(A) tail of the mRNAs susceptible to rapid degradation. As a result, acceleration of degradation of PRE-containing mRNAs by PUMs is bypassed in the absence of PABPC proteins. Intriguingly, the involvement of PABPC in PUM repression may be evolutionary conserved, as connections have been made in *Saccharomyces cerevisiae* and *Drosophila melanogaster* (45, 46).

We found that PUM and PABPC associate, based on co-immunoprecipitation assays. This observation is bolstered by in vivo data from a BioID-based proximity labeling screen, which detected interactions of PUM1 and PUM2 with PABPC1 and PABPC4 (47). At this time, the functional significance of the PUM:PABPC association is unknown. Biochemical analysis of the potential direct interaction has proven challenging for technical reasons. At this time, we cannot rule out that the PUM:PABPC association may be bridged by a mutual partner or a post-translational modification. Our data indicate that PUM associates with the RRMs of PABPC1 and does not rely on the eIF4G interaction or the MLLE motif that is recognized by PAM2 containing proteins. These results exclude the majority of known PABPC binding proteins as potential bridging factors (15). Thus, future work will be required to dissect the molecular basis of the PUM:PABPC association.

We demonstrated that the concentration of PABPC in cells controls PUM repression and mRNA stability. Increasing PABPC abundance inhibited PUM repression in a dose-dependent manner. The ability of PABPC to oppose PUM-mediated mRNA decay is the key. Over-expressed PABPC stabilizes both PUM target mRNAs and non-target mRNAs, consistent with its protection of the poly(A) tail from CCR4-NOT deadenylases. The RNA-binding activity of

PABPC is crucial; whereas, its ability to interact with eIF4G or the multitude of partners that bind to the MLLE motif was not required to counteract PUM repression. Like PABPC1, over- expression of any of the three PABPC paralogs inhibits PUM activity. In fact, PABPC5, which is composed of 4 RRMs and shares ∼65% amino acid identity with PABPC1, was capable of inhibiting the repressive activity of PUM. Together, these results indicate that the ability of PABPCs to bind and protect the poly(A) tail is their essential function in modulating PUM repression.

Based on our findings, we postulate the Goldilocks principle for PABPC function wherein PABPC levels must be “just right” to tune the response of mRNAs to regulatory sequences and trans-acting factors that control poly(A) metabolism (**Figure 10**). The overall concentration of PABPC controls mRNA decay globally and dictates responsiveness to sequence-specific regulators that, like PUMs, act through deadenylation. Too little PABPC renders mRNAs susceptible to rapid deadenylation and subsequent degradation (**Figure 10**). As a result, the action of trans-regulators to promote mRNA deadenylation is obviated, as we have demonstrated for PUMs. This Goldilocks principle is consistent with observations made by Marc Fabian’s lab using a microRNA-regulated target mRNA (29). In contrast, high concentration of PABPC promotes occupancy of poly(A) tails, thereby blocking deadenylation and degradation of mRNAs (**Figure 10**). Under this condition, the action of trans-regulators that promote deadenylation would be blocked, as we observed for PUMs.

**Figure 10.**
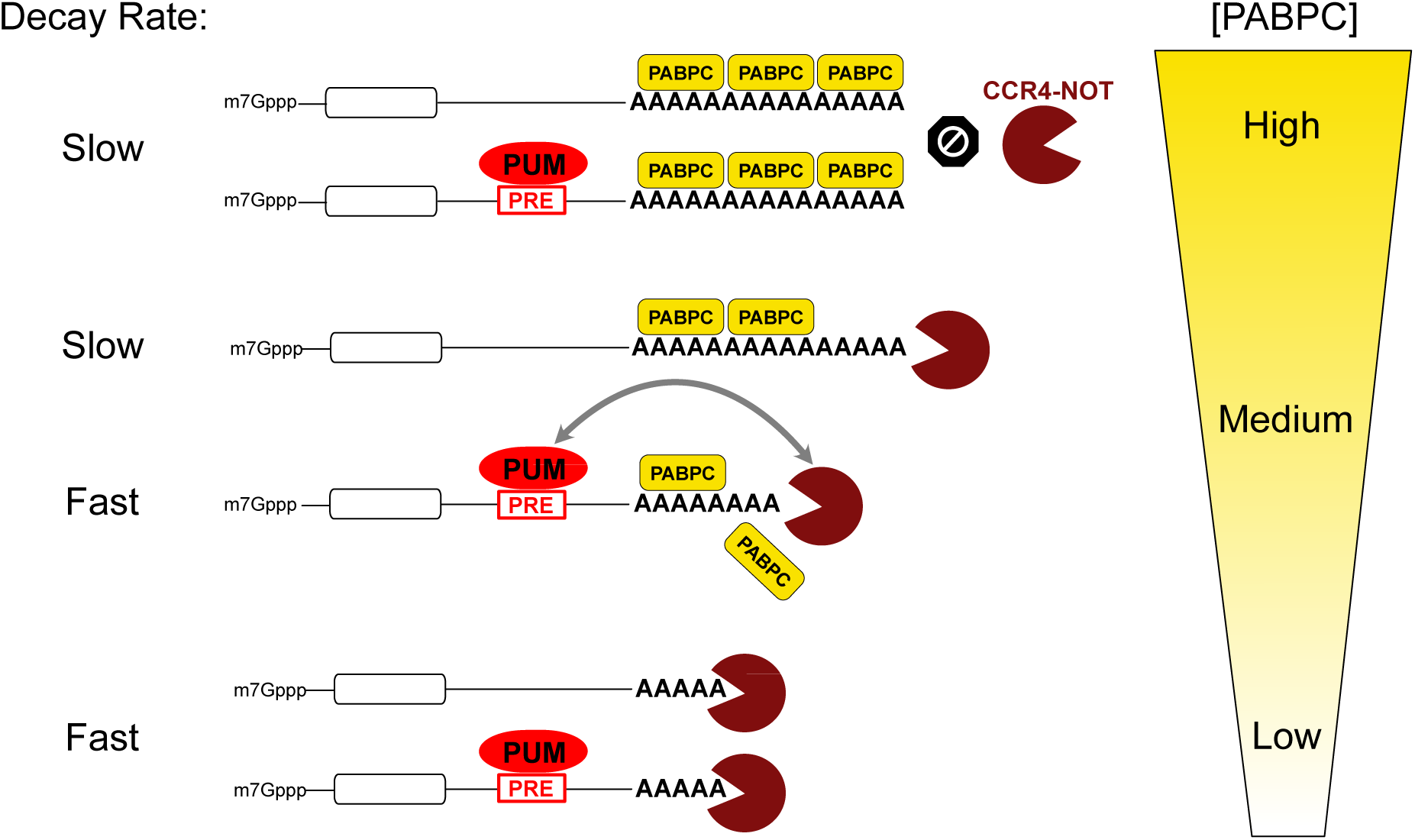
Model of the Goldilocks principle for the concentration-dependent modulation of regulated and unregulated mRNA stability by PABPC. mRNA degradation is controlled by the interplay of PABPC binding and protection of the poly(A) tail of mRNAs and the ability of deadenylases (e.g. the CCR4-NOT complex) to catalyze poly(A) shortening, leading to mRNA degradation. In contexts where PABPC concentration is intermediate “medium”, mRNA decay proceeds slowly at a basal rate, and can be accelerated by sequence specific RNA-binding repressors, such as PUM, that directly recruit the CCR4-NOT deadenylase complex. Thus, the PABPC level is “just right” to enable PUM-mediated repression, in the parlance of the eponymous fairy tail. When PABPC concentrations increase to “high” levels, deadenylation and decay of mRNAs is effectively blocked, and PUM:CCR4-NOT is no longer capable of deadenylation. When PABPC abundance decreases to “low” levels, the poly(A) tail becomes exposed and the mRNA is subject to rapid degradation, thereby bypassing the role of PUM.

The Goldilocks principle has broad implications for regulation of mRNA degradation by the growing list of sequence specific, trans-acting mRNA regulators that function through deadenylation including PUMs, Nanos, TTP, Roquin, BTG/TOB, the N6-methyl-adenosine reader YTHDF2, and microRNA-induced silencing complex (31, 65). We speculate that the Goldilocks principle may also be relevant to other deadenylation-dependent processes including codon optimality mediated mRNA decay (24, 65), P-site tRNA-mediated mRNA decay (66), and tubulin autoregulation (67).

The physiological relevance of the Goldilocks model is emphasized by evidence showing that the expression of PABPCs varies substantially across tissues, cell types, and developmental stages. The survey of PABPC expression in normal human tissues demonstrates that the abundance of PABPC1 and PABPC4 mRNA vary widely, supported by our analysis of their protein expression in human tissues. Additionally, PABPC1 expression changes during development, as dramatically demonstrated for the drastic decline of PABPC1 protein in the developing heart (68). Moreover, PABPC1 expression can be reactivated in disease states, specifically cardiac hypertrophy and certain types of cancer (69–71). Based on current knowledge, application of the Goldilocks principle should also consider the abundance of the multiple PABPC paralogs that exhibit overlapping, or in some cases unique, patterns of tissue and developmental expression (7). Together, the Goldilocks principle of PABPC function and the observed variations in PABPC levels indicates that mRNA stability and regulation are likely context specific and highly tunable.

## MATERIAL AND METHODS

### Plasmids

The oligonucleotides and plasmids used in this study are listed in **Supplemental Table S1**. The nano-luciferase (Nluc) 3xPRE and 3xPREmt reporter constructs were previously described (35–37). The Nluc reporters containing the 3′ UTRs of PUM natural targets, FZD8 and CDKN1B (**Figure 3**) were derived from constructs previously reported in Bohn et al. (36). The firefly luciferase (Fluc) plasmid pGL4.13 (Promega) was used as a transfection efficiency control. The Fluc MALAT and Nluc MALAT constructs were created by replacing the SV40 cleavage and poly-adenylation signal with the mouse MALAT1 3′ region including the triple helix, RNase P cleavage site, and Mascrna via restriction cloning (42, 43, 72). The internal A20 and A60 tracts were introduced between the PRE motifs and the MALAT1 triple helix sequence (**Figure 1**).

The Tet-Off reporter system used in **Figures 5 and 9** was based on the pTet-Off All-In-One plasmid, generously provided by Dr. Michael Kearse (73). The Tet-Off reporters contain the tetracycline-responsive PTight promoter and tTA-Advanced coding sequence from pCW57.1- MAT2A (Addgene plasmid # 100521). A 5′ UTR containing a chimeric intron and the coding region of Nluc (the same sequences used in the Nluc 3xPRE reporter assays) were subcloned into this pTet-Off plasmid. Then 6xPRE or 6xPREmt sequences were inserted into the 3′ UTR controlling the Nluc gene.

### Cell culture and transfection

Human HCT116 cells (ATCC, CCL-247) were cultured at 37 °C under 5% CO_2_ in McCoy’s 5A modified medium (Gibco, Fisher 16600082) with 10% (v/v) fetal bovine serum (Genesee 25- 514) and antibiotics (100 U/mL penicillin and 100 µg/mL streptomycin, Thermo Fisher Scientific). Transfections were performed according to the manufacturer’s instructions using 4 μl of Fugene HD (Promega, E2312) per 1 μg of plasmid DNA. Specific transfection conditions are described for each experimental method below.

### Luciferase reporter assays

For 96-well reporter assays, HCT116 cells were seeded with 5,000 cells in 100 μl of growth medium per well in a 96-well white-walled plate (Fisher, 12-566-00). After 24 hrs, the cells were transfected with 5 ng Fluc reporter plasmid, 10 ng of Nluc reporter plasmid, and 85 ng pF5A empty vector control or effector protein expression vector (100 ng total transfected DNA per well). For reporter assays with PABPC titrations as in **Figure 7**, the cells were transfected with 5 ng Fluc MALAT reporter plasmid, 10 ng of Nluc reporter plasmid, and the indicated mass of PABPC recombinant protein plasmid. The amount of transfected PABPC plasmid was balanced with the pF5A control vector to maintain a total of 85 ng per sample. The transfected cells were grown for another 48 hrs and luciferase values were measured using the Dual-Glo Assay system (Promega) and a GloMax Discover luminometer.

Reporter data was analyzed as described previously (35–37). A relative response ratio (RRR) was calculated by normalizing the Nluc luciferase activity (measured in relative light units, RLU) against the corresponding Fluc RLU for each sample. These RRR values were then used to calculate the fold change between the Nluc 3xPRE relative to the Nluc 3xPREmt to measure repression by endogenous PUMs. All reporter assays were performed in three independent experiments, each of which had 3 biological replicates, for a total of nine replicates. The data is reported as mean log(2) fold change along with the standard deviation. The statistical significance of comparisons was calculated using an ordinary one-way ANOVA and Tukey test for multiple comparisons. The resulting data, statistics, and number and type of replicates are reported in the figure legends and in **Supplemental Table S2**.

### RNA interference

For RNAi knockdown experiments, HCT116 cells were first seeded at 200,000 cells in 2 ml per well in a 6-well plate (USA Scientific CC7682-7506). After 24 hrs, the cells were transfected with 25 nM final concentration of On-Target Plus Smartpool siRNAs (Horizon Discovery) for the indicated gene of interest, listed in **Supplemental Table S1**. On-Target Plus non-targeting control siRNAs (NTC) were used as the negative control for all RNAi experiments. For the PABPC1 RNAi knockdown, 50 nM siRNAs per transfection was used, and a second siRNA transfection at 48 hrs after seeding was necessary for efficient depletion. For knockdowns in reporter assays, the reporter constructs were transfected 24 hrs after the last siRNA treatment. The cells were transfected with 0.5 μg Fluc MALAT reporter plasmid, 1.25 μg of Nluc reporter plasmid, and 1.25 μg pF5A empty vector control or effector protein expression vector (3 μg total DNA). The cells were harvested 48 hrs after the reporter transfection and re-seeded at 50,000 cells per well into a 96-well plate for luciferase reporter assays described above. Each RNAi knockdown was verified by western blot analysis.

To measure changes in endogenous PUM target mRNAs (**Figure 6**), siRNAs targeting PUM1 and PUM2 were transfected into the HCT116 AtAFB2 cell line. These cells were seeded at 200,000 per well in 6-well plates and transfected 24 hours later with either 25 nM NTC siRNAs or 12.5 nM each of PUM1&2 siRNAs, totaling 25 nM.

### PABPC1 IAA-induced degradation

The HCT116 PABPC1-AID cell line (sC278-C2) was generously provided by Drs. Kehui Xiang and David Bartel and was described in Xiang and Bartel, eLife, 2021 (30). These cells were seeded at 400,000 cells per well in a 6-well plate. After 24 hr incubation, 1 µg/ml of doxycycline (Sigma, D9891) in dimethyl sulfoxide (DMSO) was added to the cells to induce OsTiR expression along with 0.2 mM auxinole (Aobious, AOB8812), an inhibitor of OsTIR1 used to prevent “leaky” degradation. After 30 mins, luciferase reporter plasmids were transfected with 0.5 μg Fluc MALAT plasmid, 1.25 μg of Nluc reporter plasmid, and 1.25 μg pF5A empty vector control (3 μg total DNA). After 6 hrs, fresh media containing 1 µg/ml of doxycycline and 0.5 mM of indole-3-acetic acid (IAA, Goldbio, I-110-25) were added to the test samples. The control samples were incubated with fresh media with the equivalent volume of DMSO as a vehicle-only control. Twenty-four hours after reporter transfection, the cells were trypsinized and reseeded at 50,000 cells per well into a 96-well plate for luciferase reporter assays as described above. The remaining cells were collected by centrifugation to analyze by western blotting.

For PABPC4 RNAi and PABPC1-AID mediated double knockdowns, 400,000 HCT116 PABPC1-AID cells per well were seeded in a 6-well plate. 24 hrs post-seeding, the cells were transfected with siRNAs as detailed above. After another 24 hr incubation, 1 µg/ml of doxycycline and 0.2 mM auxinole was added to each well (including all controls). After 30 mins, luciferase reporter plasmids were transfected. The cells were incubated for 6 hrs and fresh media containing 1 µg/ml of doxycycline and 0.5 mM IAA was added to the test samples. 24 hrs after the reporter transfection, the cells were trypsinized and re-seeded at 50,000 cells per well into a 96-well plate for luciferase reporter assays as described above. The remaining cells were split, collected by centrifugation, and analyzed by western blotting.

To analyze endogenous gene expression in response to PABPC1-AID degradation and PABPC4 RNAi depletion (**Figure 6**), HCT116 PABPC1-AID cells were seeded at 400,000 cells per well in 6-well plates. After 24 hours, the cells were transfected with either 25 nM siNTC or 25 nM siPABPC4. Twenty-four hours after siRNA transfection, 1 µg/mL doxycycline (w/v in DMSO) was added to induce OsTIR expression, along with 0.2 mM auxinole in all samples. Six hours later, the medium was replaced with fresh medium containing 1 µg/mL doxycycline and 0.5 mM IAA for test samples, while control samples received medium with an equivalent volume of DMSO as a vehicle control. All conditions were performed with three biological replicates. After a total of 18 hours of IAA treatment to deplete PABPC1-AID and 72 hours of RNAi to deplete PABPC4, cells were harvested for analysis of RNA and protein. Depletion of PABPC1 and PABPC4 was verified by western blot analysis. The RNA collected was used for RTqPCR to assess the responses of endogenous genes, as described below.

### Immunoprecipitation assays

For co-immunoprecipitation (co-IP) of endogenous proteins (**Figure 3**), HCT116 cells were seeded at 1x10^6^ cells in 10 mLs in a T75 flask and grown until confluent. First, the antibody was pre-bound to magnetic beads. For each sample, 20 µL of magnetic protein A surebeads (Biorad) were equilibrated 3 times 250 µL 1x phosphate buffered saline (PBS) pH 7.4 (Thermo Fisher Scientific) + 0.1% Tween-20 and incubated in 100 µl 1xPBS + 0.1% Tween-20 + 2 µg PUM1 antibody (Bethyl labs, A302-576A) overnight, rotating at 4°C. Next, cell extracts were prepared. The HCT116 cells were collected by centrifugation at 500 x g for 5 mins. and lysed by cell disruption in 400 µl of lysis buffer (50 mM Tris-HCl (pH 8.0), 500 mM NaCl, 0.5% Triton X100, 1 mM EDTA) with 2x complete protease inhibitor cocktail (Roche, 11836153001). Cell debris was removed by centrifugation at 10,000 x g for 10 mins and the supernatant was then passed through a 0.45-micron filter at 4,000 x g for 3 mins. The protein concentration of the extract was measured using the DC-Lowry assay (BioRad). Next, 500 µg of total cellular protein in 200 µl of lysis buffer with 4 μg of RNase A (Promega) and 40 units of RNase ONE (Promega) was added to the antibody-bound protein A surebeads and incubated for 2 hrs, rotating at 4°C . The beads were washed 6 times with 500 µl lysis buffer and eluted in 25 µl SDS-PAGE loading dye (80 mM Tris-HCl pH 6.8, 10% glycerol, 2% SDS, 100 mM DTT, 0.2% Bromphenol Blue, 100 mM 2-Mercaptoethanol) at 85°C for 10 min. The supernatant was separated from the beads using a magnet and 50% of the immunoprecipitate was analyzed by western blot along with 2% of the cell lysate (i.e. input).

Co-immunoprecipitation of proteins expressed from transfected plasmids (**Figure 3**), HT, HT- PABPC1 full-length, HT-PABPC1 eIF4Gmt, HT-PABPC1 MLLEmt, HT-PABPC1 MLLE domain, and HT-PABPC1 RRM1-4 domain were performed in the same manner with the following changes. Cells were seeded in a 6-well plate at 200,000 cells in 2 ml per well. After 24 hrs, unless indicated in the figure legend, the cells were transfected with 1.5 μg effector protein expression vector and 1.5 μg of pF5A empty vector control (3 μg total DNA). 48 hrs post-transfection, two wells were harvested for each co-IP sample. The PUM1 antibody-bound beads were incubated with 300 µg of total protein lysate, per sample.

For RNA co-immunoprecipitation assays (**Figure 4 and Figure 8**), HCT116 cells were seeded at 200,000 cells per well in a 6-well plate. For experiments wherein PABPC1 was immunoprecipitated, cells were transfected with 1.5 μg of either the 3xPRE or 3xPREmt Nluc reporter plasmid and 1.5 μg of pF5A empty vector control (3 μg total DNA). For immunoprecipitation of PUM1 protein, cells were transfected with 1.5 μg of 3xPRE reporter plasmid and 1.5 μg of either HT control or the HT-PABPC1 effector expression plasmid, as indicated in the figure. Two wells were harvested per sample. For each sample, 8.3 µl of protein A conjugated dynabeads (Thermo Fisher Scientific, 10001D) were washed 3 times with 250 µl 1xPBS + 0.1% Tween-20 and incubated in 100 µl of 1xPBS + 0.1% Tween-20 and 2 µg PABPC1 antibody (Abcam, ab6125) or PUM1 antibody (Bethyl labs, A302-576A) overnight, rotating at 4°C. The HCT116 cells were collected by centrifugation at 500 x g for 5 mins and lysed by cell disruption in 400 µl of lysis buffer (50 mM Tris-HCl (pH 8.0), 500 mM NaCl, 0.5% Triton-X100, 1 mM EDTA, RNasin (Promega) with 2x complete protease inhibitor cocktail (Roche, 11836153001). Cell debris was removed by centrifugation at 10,000 x g for 10 mins and the supernatant was then passed through a 0.45-micron filter at 4,000 g for 3 mins. The protein concentration of the extract was measured using the DC-Lowry assay (BioRad). For each sample, 10 µl of lysate was set aside as the “input” sample. Next, 300 µg of total protein in 200 µl of lysis buffer was added to the protein A dynabeads and incubated for 2 hrs with end over end rotation at 4°C. The supernatant was removed from the beads, which were subsequently washed 6x with 500 µl of lysis buffer without RNasin. The beads were suspended in a final volume of 20 µl, and 5 µl of which was reserved for western blot analysis (described below). RNA was purified from the input sample and the remaining beads using the RNA clean and concentrator 5 kit (Zymo, R1013) following the manufacturer’s instructions, including the on-bead DNase I digestion. Purified RNA was eluted in a final volume of 10 µl ddH_2_0. The entire RNA sample was analyzed by northern blotting, along with 1% of the RNA purified from the input sample. Note that ethidium bromide was not included in the sample loading buffer during the formaldehyde-agarose gel electrophoresis, as it can alter the electrophoretic mobility of the RNA.

### PABPC1-AID cells

For the mRNA decay experiments (**Figure 5**), we created an auxin inducible degron tagged PABPC1 cell line based on the AtAFB2 system that does not require induction using doxycycline. To generate the HCT116 PABPC1-AID AtAFB2 cell line, the AID degron tag (miniIAA7) (60) with a 3xFlag epitope tag and flexible linker was introduced at the C-terminus of the PABPC1 coding region using Cas9-mediated genome engineering in the HCT116 AtAFB2 parental cell line (generously provided by Dr. Eric Wagner, Univ of Rochester). Homology arms of 500 nt flanking the stop codon of PABPC1 with a linker-miniIAA7-3xFlag were ordered from Twist biosciences and cloned into a donor plasmid (gift from Wagner lab) derived from a pcDNA3 vector as described here (74). The PAM region targeted by Cas9 within the donor plasmid was mutated. The Cas9 single guide RNA was cloned into pX459-V2.0 plasmid (Addgene plasmid # 62988) by oligo cloning. The HDR donor plasmid and sgRNA were co-transfected into the HCT116 AtAFB2 cells. Cells transfected with the donor template were selected using 18 µg/ml blasticidin (Thermo Fisher Scientific, A1113903) for 2 days. Clonal isolation was performed by limiting dilution in 96-well plates. The clones were expanded and genotyped by PCR (Phire Tissue Direct PCR Master Mix, Thermo Fisher Scientific, F170L) and western blotting (**Figure S3B and S3C**). The clone used in this study was confirmed to be homozygous by a single shifted band in the genomic PCR and DNA sequencing of the PCR product.

### Transcription shut-off and mRNA decay assays

To measure RNA decay rates of the Nluc mRNAs as in **Figure 5**, HCT116 PABPC1-AID AtAFB2 cells were seeded at 400,000 cells in 2 ml per well. 24 hrs post seeding, cells were transfected with either non-targeting control siRNAs or PABPC4 siRNAs as described above. 48 hrs post-seeding, the cells were transfected with 750 ng pTet-Off All-In-One plasmids containing Nluc with 6xPRE or 6xPREmt in the 3′ UTR, 1.75 μg pF5A empty vector control, and 0.5 μg Fluc MALAT1 reporter plasmid. Directly after transfection, 25 ng/ml of doxycycline was added to turn off transcription in all conditions. 0.5 mM of IAA was added to the PABPC4 siRNA treated wells to begin PABPC1 depletion and equivalent volume of DMSO vehicle control was added to the NTC control wells. 24 hrs later, the cells were washed with 1xPBS and fresh media without doxycycline was added and the cells were incubated at 37° C for 2 hrs to induce transcription of the Nluc mRNA. 0.5 mM of IAA was included in the media for the PABPC1&C4 samples. After 2 hrs, transcription was shut off by adding fresh media containing 2 μg/mL doxycycline to all the conditions. Cells were then harvested at the 0, 1, 2, 4, 6, and 8 hr timepoints and RNA was extracted and analyzed by Chemi-Northern blotting as described below. The resulting half-lives were determined from first order exponential decay trend lines calculated by nonlinear regression using GraphPad Prism (76).

### PABPC1 over-expression for northern blot analysis

The effect of HT-PABPC1 over-expression on Nluc mRNA levels was measured in HCT116 cells in **Figure 9**. Cells were seeded at 400,000 cells per well in 2 mL of growth medium in a 6- well plate. Twenty-four hours after seeding, cells were transfected using Fugene HD with 0.125 µg of either Nluc 6xPRE or Nluc 6xPREmt reporter plasmid, 1.2 µg of pFN21K HT PABPC1 or 0.5 µg of HaloTag plasmid, and 0.5 µg of Fluc MALAT plasmid. The total mass of transfected DNA was balanced using pF5A empty vector for a total of 3 µg of DNA per well. Twenty-four hours post-transfection, RNA and protein were harvested and analyzed by northern and western blot, respectively. For the Nluc mRNA marker with the poly(A) tail removed (A0), a control RNA sample was treated with oligo-dT15 (Promega) and RNase H (New England Biolabs) as previously described (77). This experiment included three biological replicates per condition.

### RNA purification

HCT116 cells were harvested by washing twice with PBS pH 7.4, followed by trypsinization with TrypLE (Thermo Fisher Scientific) for 5 mins, and centrifugation at 500 g for 5 mins. RNA was purified from the cell pellet using the SimplyRNA cells kit (Promega, AS1390) and Maxwell RSC instrument following the manufacturer’s protocol. For the on-bead DNase digestion, the amount of DNase I (Promega, Z3585) was doubled (10 μL total) to ensure removal of genomic and plasmid DNA. Purified RNA was eluted in 40 μL of nuclease-free water and quantified using a NanoDrop spectrophotometer (Thermo Scientific).

### Northern blotting

Northern blotting was performed using the chemi-northern method as previously described (78). The anti-sense probes used in this study are listed in **Supplemental Table S1.** Briefly, 1 µg of RNA was used in the time course and steady-state overexpression experiments (Figure 9), whereas for all other northern blots, 2.5 µg of purified total RNA per sample, or 10 µl of RIP samples were prepared in a final volume of 24 µL containing final concentrations of northern RNA sample buffer (0.04 μg/μL ethidium bromide, 23% formamide, 3% formaldehyde, 4.6 mM 3-(N-Morpholino)propanesulfonic acid hemisodium salt (MOPS) pH 7.0, 1.1 mM sodium acetate, and 0.2 mM EDTA), and northern RNA loading dye (2.1% glycerol, 4.2 mM EDTA and 0.01% (w/v) of Bromophenol Blue and Xylene Cyanol) along with RNA molecular weight markers (Promega, G3191). The samples were heated at 70 °C for 10 mins and separated on a denaturing formaldehyde-agarose gel (1% (w/v) agarose gel containing 1x MOPS buffer (20 mM MOPS pH 7.0, 5 mM sodium acetate, and 1 mM EDTA with 1.48% formaldehyde) in 1x MOPs running buffer at 95 V for 1.5 hrs. The RNA was transferred to positively charged nylon membrane (Immobilon-Ny+, Millipore, INYC00010) by downward capillary transfer overnight in 20x SSC buffer containing 3 M NaCl and 300 mM sodium citrate (pH 7.0). After transfer, the blot was exposed to UV (120 J/cm2, λ = 254 nm) in a CL-1000 crosslinker (UVP). To detect the RNA marker, its lane was removed from blot and visualized by staining with 0.25% w/v methylene blue in ddH_2_O for 5 mins, followed by washing in ddH_2_O until marker bands were clearly visible. The blot was incubated with rotation in 10 mL pre-warmed Ultrahyb Ultrasensitive hybridization buffer (Thermo Fisher Scientific, AM8670) at 68 °C for 1 hr for RNA-based probes (Nluc) or 10 mL of pre-warmed Ultrahyb oligo hybridization buffer (Thermo Fisher Scientific, AM8663) at 42 °C for 1 hr for DNA oligo probes (18S). The Nluc riboprobe was hybridized at 50 ng/ml and the 18S DNA oligo probe was hybridized at 5 nM. Hybridization proceeded for 12 or more hours at 68 °C (riboprobe) or 42 °C (oligo probe). After probe hybridization, the buffer was removed from the bottle and then the blot was washed twice with 2xSSC and 0.1% SDS and then twice with 0.1xSSC and 0.1% SDS. Each wash was performed at 68 °C (RNA probe) or 42 °C (DNA oligo probe) using 25 mL of the stated buffer for 15 minutes each in a rotisserie hybridization oven. The Chemiluminescent Nucleic Acid Detection Module (Thermo Fisher Scientific, 89880) was used to detect the biotinylated probe on the blot with streptavidin-HRP and enhanced chemiluminescence using the manufacturer’s instructions and supplied reagents. The blot was visualized using a chemiluminescence imaging system (Azure 300 Chemiluminescent Imager, Azure Biosystems).

### Western blotting

For western analysis of the reporter assays, cells were lysed by cell disruption in 100 µL of RIPA buffer (25 mM Tris-HCl pH 7.6, 150 mM NaCl, 1% NP-40, 1% sodium deoxycholate, 0.1% SDS) supplemented with 2x complete protease inhibitor cocktail (Roche). The protein concentration was determined using the DC-Lowry assay (BioRad). Unless indicated, 10 µg of total protein lysate per sample was prepared in SDS-PAGE loading dye (80 mM Tris-HCl pH 6.8, 10% glycerol, 2% SDS, 100 mM DTT, 0.2% bromphenol blue, 100 mM 2-mercaptoethanol), heated to 85°C for 10 mins, and separated by SDS-PAGE on a 4-20% gradient polyacrylamide gel (Bio-Rad). The proteins were then transferred to a PVDF membrane (Immobilon P, Millipore) at 60V constant for 1.5 hrs at 4 °C. The membrane was blocked for 1 hr in 5% w/v nonfat powdered dry milk, 1x PBS, and 0.1% Tween-20 (PBST) and incubated in the indicated primary antibody diluted to 1:1,000 in PBST overnight at 4 °C. The membrane was washed 3 times, for 10 mins each, with PBST and incubated with the appropriate HRP secondary antibody diluted 1:10,000 in PBST. The membrane was washed 3 more times with PBST and visualized by enhanced chemiluminescence using Immobilon Western Chemiluminescent HRP Substrate (Millipore). The antibodies and dilutions used in this study are detailed in **Supplemental Table S1.**

### Analysis and quantitation of northern and western blots

The chemi-northern and western blots were analyzed using AzureSpot Pro analysis software (Azure Biosystems). Individual lanes were identified automatically and adjusted manually as needed. Band signals were detected using fixed band width and the background was corrected using the rolling ball method with the radius set to 20%.

For northern blot quantitation of PUM repressive activity, the Nluc mRNA signal intensities were normalized to their respective loading controls, 18S rRNA, on the corresponding blots. The resulting ratios were then used to calculate the fold change of the PUM-regulated Nluc 3xPRE reporter relative to the unregulated mutant reporter, Nluc 3xPREmt. Measurements were obtained for three biological replicates for each experimental condition. For the PABPC1 and PUM1 RNA coimmunoprecipitation assays, the Nluc mRNA signal intensities of the immunoprecipitate samples were normalized to their respective inputs. The fold enrichment of the normalized PABPC1 RIPs was then calculated relative to the IgG negative control samples. Measurements were obtained for three biological replicates for each experimental condition.

### Reverse transcription and quantitative polymerase chain reaction

The RTqPCR parameters, including the primer sequences, amplification efficiencies, and amplicon sizes, following the standard MIQE guidelines are reported in **Supplemental Table S1**. Gene expression of ITGA2, SMPDL3A, and GAPDH mRNAs and MALAT1 ncRNAs was analyzed in PUM1&2 and PABPC1&C4 knockdown conditions (**Figure 6**). RNA was isolated using the Maxwell system, and 1 µg of RNA was reverse transcribed with the LunaScript RT kit (New England Biolabs) and 1 µL of random hexamer oligonucleotides (Promega) per reaction. Reverse transcription was performed at 25°C for 2 min, 55°C for 10 min, and 95°C for 1 min.

Quantitative PCR was performed using 50 ng of cDNA for ITGA2, MALAT1, GAPDH, or 0.4 ng of cDNA for 18S rRNA, or 83.3 ng of cDNA for SMPDL3A. Each reaction contained 0.25 µM of each primer and Luna Universal qPCR Master Mix (New England Biolabs), following the manufacturer’s protocol. As negative control, a no-RT reaction was analyzed for each primer set. The cycling conditions were: (i) 95°C for 2 min, (ii) 95°C for 15 sec, and (iii) 60°C for 1 min. Steps ii-iii were repeated for a total of 40 cycles. A melt curve was generated over a range of 65°C to 95°C. Products were analyzed by gel electrophoresis to verify the correct amplicon size. Quantification cycle (Cq) values were obtained using CFX Manager software (Bio-Rad).

Relative expression was calculated using the Livak method (2−ΔΔCq), with normalization to 18S rRNA and fold change calculations made relative to the corresponding non-depleted condition. All experiments were performed with three biological replicates, each with a corresponding no-RT control.

### Cell Count and Viability Measurements

HCT116 cells collected from reporter assays under PABPC1 and PABPC4 depletion conditions versus NTC, -IAA conditions were counted and assessed for cell viability by trypan blue staining (Thermo Fisher) using the Countess Automated Cell Counter (Thermo Fisher). These measurements were taken from three independent experiments with a total of twelve replicates per condition (**Figure 3**).

## Supporting Information

**Supplemental Figure S1. Comparison of Nluc luciferase expression from reporter mRNAs shown in Figure 1**.

**Supplemental Figure S2. Depletion of PABPC1 reduces PUM-mediated repression.**

**Supplemental Figure S3. PABPC1&4 depletion destabilizes mRNAs.**

**Supplemental Figure S4. over-expression of cytoplasmic PABP homologs alleviates PUM-repression.**

**Supplemental Figure S5. PABPC1 over-expression stabilizes mRNAs.**

**Supplemental Figure S6. PABPC1&4 mRNA expression levels across normal human tissues.**

**Supplemental Figure S7. PABPC protein expression across normal human tissues.**

**Supplemental Table S1. MIQE guidelines, oligonucleotides, plasmids, and antibodies**

**Supplemental Table S2. Data and statistics**

## Acknowledgements

We thank members of the Goldstrohm Lab, including Dr. Robert Connacher, for their feedback and critiques on this manuscript. We thank Dr. Eric Wagner and Dr. Michael Kearse for sharing reagents and Dr. Eugene Valkov and Dr. Zachary Campbell for comments and suggestions on the manuscript.

## Author Contributions

K.M.: conceptualization, investigation, methodology, visualization, formal analysis, writing, review, editing

C.H.P.: investigation, methodology, visualization, formal analysis, writing, review, editing E.D.: investigation, methodology, visualization, formal analysis, review, editing

J.P.: investigation, formal analysis

A.G.: funding acquisition, project administration, supervision, conceptualization, visualization, formal analysis, writing, review, editing

## Funding and additional information

This research was supported by National Institutes of Health grant R01 GM150468 to A.C.G. K.M.M was supported in part by the American Cancer Society Postdoctoral Fellowship 134102- PF-19-128-01-DDC. E.B.D was supported by University of Minnesota’s Targets of Cancer Training Program, NIH grant T32 CA009138. The content is solely the responsibility of the authors and does not necessarily represent the official views of the National Institutes of Health.

## Data Availability

All RTqPCR data and MIQE information is reported in Supplemental Table S1. All siRNA and oligonucleotide data is reported in Supplemental Table S1. All data, statistics, and number and type of replicates are reported in the figures, legends, and in Supplemental Table S2.

## References

1. Wells, S.E., Hillner, P.E., Vale, R.D. and Sachs, A.B. (1998) Circularization of mRNA by eukaryotic translation initiation factors. Mol. Cell, 2, 135–140.

2. Gallie, D.R. (1998) A tale of two termini: a functional interaction between the termini of an mRNA is a prerequisite for efficient translation initiation. Gene, 216, 1–11.

3. Gallie, D.R. (1991) The cap and poly(A) tail function synergistically to regulate mRNA translational efficiency. Genes Dev., 5, 2108–2116.

4. Sonenberg, N. and Hinnebusch, A.G. (2009) Regulation of translation initiation in eukaryotes: mechanisms and biological targets. Cell, 136, 731–745.

5. Kessler, S.H. and Sachs, A.B. (1998) RNA recognition motif 2 of yeast Pab1p is required for its functional interaction with eukaryotic translation initiation factor 4G. Mol. Cell. Biol., 18, 51–57.

6. Gebauer, F. and Hentze, M.W. (2004) Molecular mechanisms of translational control. Nat. Rev. Mol. Cell Biol., 5, 827–835.

7. Smith, R.W.P., Blee, T.K.P. and Gray, N.K. (2014) Poly(A)-binding proteins are required for diverse biological processes in metazoans. Biochem. Soc. Trans., 42, 1229–1237.

8. Guzeloglu-Kayisli, O., Pauli, S., Demir, H., Lalioti, M.D., Sakkas, D. and Seli, E. (2008) Identification and characterization of human embryonic poly(A) binding protein (EPAB). Mol. Hum. Reprod., 14, 581–588.

9. Yang, H., Duckett, C.S. and Lindsten, T. (1995) iPABP, an inducible poly(A)-binding protein detected in activated human T cells. Mol. Cell. Biol., 15, 6770–6776.

10. Kini, H.K., Kong, J. and Liebhaber, S.A. (2014) Cytoplasmic poly(A) binding protein C4 serves a critical role in erythroid differentiation. Mol. Cell. Biol., 34, 1300–1309.

11. Kleene, K.C., Mulligan, E., Steiger, D., Donohue, K. and Mastrangelo, M.A. (1998) The mouse gene encoding the testis-specific isoform of Poly(A) binding protein (Pabp2) is an expressed retroposon: intimations that gene expression in spermatogenic cells facilitates the creation of new genes. J. Mol. Evol., 47, 275–281.

12. Blanco, P., Sargent, C.A., Boucher, C.A., Howell, G., Ross, M. and Affara, N.A. (2001) A novel poly(A)-binding protein gene (PABPC5) maps to an X-specific subinterval in the Xq21.3/Yp11.2 homology block of the human sex chromosomes. Genomics, 74, 1–11.

13. Féral, C., Guellaën, G. and Pawlak, A. (2001) Human testis expresses a specific poly(A)- binding protein. Nucleic Acids Res., 29, 1872–1883.

14. Goldstrohm, A.C. and Wickens, M. (2008) Multifunctional deadenylase complexes diversify mRNA control. Nat. Rev. Mol. Cell Biol., 9, 337–344.

15. Passmore, L.A. and Coller, J. (2022) Roles of mRNA poly(A) tails in regulation of eukaryotic gene expression. Nat. Rev. Mol. Cell Biol., 23, 93–106.

16. Dowdle, M.E. and Lykke-Andersen, J. (2025) Cytoplasmic mRNA decay and quality control machineries in eukaryotes. Nat. Rev. Genet., 26, 463–478.

17. Schäfer, I.B., Yamashita, M., Schuller, J.M., Schüssler, S., Reichelt, P., Strauss, M. and Conti, E. (2019) Molecular Basis for poly(A) RNP Architecture and Recognition by the Pan2- Pan3 Deadenylase. Cell, 177, 1619–1631.e21.

18. Uchida, N., Hoshino, S.-I. and Katada, T. (2004) Identification of a human cytoplasmic poly(A) nuclease complex stimulated by poly(A)-binding protein. J. Biol. Chem., 279, 1383–1391.

19. Wilson, T. and Treisman, R. (1988) Removal of poly(A) and consequent degradation of c-fos mRNA facilitated by 3’ AU-rich sequences. Nature, 336, 396–399.

20. Swartwout, S.G. and Kinniburgh, A.J. (1989) c-myc RNA degradation in growing and differentiating cells: possible alternate pathways. Mol. Cell. Biol., 9, 288–295.

21. Brewer, G. and Ross, J. (1988) Poly(A) shortening and degradation of the 3’ A+U-rich sequences of human c-myc mRNA in a cell-free system. Mol. Cell. Biol., 8, 1697–1708.

22. Bernstein, P., Peltz, S.W. and Ross, J. (1989) The poly(A)-poly(A)-binding protein complex is a major determinant of mRNA stability in vitro. Mol. Cell. Biol., 9, 659–670.

23. Simón, E. and Séraphin, B. (2007) A specific role for the C-terminal region of the Poly(A)- binding protein in mRNA decay. Nucleic Acids Res., 35, 6017–6028.

24. Tucker, M., Staples, R.R., Valencia-Sanchez, M.A., Muhlrad, D. and Parker, R. (2002) Ccr4p is the catalytic subunit of a Ccr4p/Pop2p/Notp mRNA deadenylase complex in Saccharomyces cerevisiae. EMBO J., 21, 1427–1436.

25. Viswanathan, P., Ohn, T., Chiang, Y.-C., Chen, J. and Denis, C.L. (2004) Mouse CAF1 can function as a processive deadenylase/3’-5’-exonuclease in vitro but in yeast the deadenylase function of CAF1 is not required for mRNA poly(A) removal. J. Biol. Chem., 279, 23988–23995.

26. Yi, H., Park, J., Ha, M., Lim, J., Chang, H. and Kim, V.N. (2018) PABP Cooperates with the CCR4-NOT Complex to Promote mRNA Deadenylation and Block Precocious Decay. Mol. Cell, 70, 1081–1088.e5.

27. Webster, M.W., Chen, Y.-H., Stowell, J.A.W., Alhusaini, N., Sweet, T., Graveley, B.R., Coller, J. and Passmore, L.A. (2018) mRNA Deadenylation Is Coupled to Translation Rates by the Differential Activities of Ccr4-Not Nucleases. Mol. Cell, 70, 1089–1100.e8.

28. Yamashita, A., Chang, T.-C., Yamashita, Y., Zhu, W., Zhong, Z., Chen, C.-Y.A. and Shyu, A.-B. (2005) Concerted action of poly(A) nucleases and decapping enzyme in mammalian mRNA turnover. Nat. Struct. Mol. Biol., 12, 1054–1063.

29. Kajjo, S., Sharma, S., Chen, S., Brothers, W.R., Cott, M., Hasaj, B., Jovanovic, P., Larsson, O. and Fabian, M.R. (2022) PABP prevents the untimely decay of select mRNA populations in human cells. EMBO J., 41, e108650.

30. Xiang, K. and Bartel, D.P. (2021) The molecular basis of coupling between poly(A)-tail length and translational efficiency. eLife, 10, e66493.

31. Raisch, T. and Valkov, E. (2022) Regulation of the multisubunit CCR4-NOT deadenylase in the initiation of mRNA degradation. Curr. Opin. Struct. Biol., 77, 102460.

32. Wang, X., McLachlan, J., Zamore, P.D. and Hall, T.M.T. (2002) Modular recognition of RNA by a human pumilio-homology domain. Cell, 110, 501–512.

33. Goldstrohm, A.C., Hall, T.M.T. and McKenney, K.M. (2018) Post-transcriptional Regulatory Functions of Mammalian Pumilio Proteins. Trends Genet., 34, 972–990.

34. Eisen, T.J., Eichhorn, S.W., Subtelny, A.O., Lin, K.S., McGeary, S.E., Gupta, S. and Bartel, D.P. (2020) The Dynamics of Cytoplasmic mRNA Metabolism. Mol. Cell, 77, 786–799.e10.

35. Van Etten, J., Schagat, T.L., Hrit, J., Weidmann, C.A., Brumbaugh, J., Coon, J.J. and Goldstrohm, A.C. (2012) Human Pumilio Proteins Recruit Multiple Deadenylases to Efficiently Repress Messenger RNAs. J. Biol. Chem., 287, 36370–36383.

36. Bohn, J.A., Van Etten, J.L., Schagat, T.L., Bowman, B.M., McEachin, R.C., Freddolino, L. and Goldstrohm, A.C. (2018) Identification of diverse target RNAs that are functionally regulated by human Pumilio proteins. Nucleic Acids Res., 46, 362–386.

37. Enwerem, I.I.I., Elrod, N.D., Chang, C.-T., Lin, A., Ji, P., Bohn, J.A., Levdansky, Y., Wagner, E.J., Valkov, E. and Goldstrohm, A.C. (2021) Human Pumilio proteins directly bind the CCR4-NOT deadenylase complex to regulate the transcriptome. RNA, 27, 445–464.

38. Gennarino, V.A., Singh, R.K., White, J.J., De Maio, A., Han, K., Kim, J.-Y., Jafar-Nejad, P., di Ronza, A., Kang, H., Sayegh, L.S., et al. (2015) Pumilio1 haploinsufficiency leads to SCA1-like neurodegeneration by increasing wild-type Ataxin1 levels. Cell, 160, 1087–1098.

39. Gennarino, V.A., Palmer, E.E., McDonell, L.M., Wang, L., Adamski, C.J., Koire, A., See, L., Chen, C.-A., Schaaf, C.P., Rosenfeld, J.A., et al. (2018) A Mild PUM1 Mutation Is Associated with Adult-Onset Ataxia, whereas Haploinsufficiency Causes Developmental Delay and Seizures. Cell, 172, 924–936.e11.

40. Gong, Y., Liu, Z., Yuan, Y., Yang, Z., Zhang, J., Lu, Q., Wang, W., Fang, C., Lin, H. and Liu, S. (2022) PUMILIO proteins promote colorectal cancer growth via suppressing p21. Nat. Commun., 13, 1627.

41. Naudin, C., Hattabi, A., Michelet, F., Miri-Nezhad, A., Benyoucef, A., Pflumio, F., Guillonneau, F., Fichelson, S., Vigon, I., Dusanter-Fourt, I., et al. (2017) PUMILIO/FOXP1 signaling drives expansion of hematopoietic stem/progenitor and leukemia cells. Blood, 129, 2493–2506.

42. Wilusz, J.E. (2016) Long noncoding RNAs: Re-writing dogmas of RNA processing and stability. Biochim. Biophys. Acta BBA - Gene Regul. Mech., 1859, 128–138.

43. Brown, J.A., Bulkley, D., Wang, J., Valenstein, M.L., Yario, T.A., Steitz, T.A. and Steitz, J.A. (2014) Structural insights into the stabilization of MALAT1 noncoding RNA by a bipartite triple helix. Nat. Struct. Mol. Biol., 21, 633–640.

44. Wilusz, J.E., JnBaptiste, C.K., Lu, L.Y., Kuhn, C.-D., Joshua-Tor, L. and Sharp, P.A. (2012) A triple helix stabilizes the 3’ ends of long noncoding RNAs that lack poly(A) tails. Genes Dev., 26, 2392–2407.

45. Chritton, J.J. and Wickens, M. (2011) A role for the poly(A)-binding protein Pab1p in PUF protein-mediated repression. J. Biol. Chem., 286, 33268–33278.

46. Weidmann, C.A., Raynard, N.A., Blewett, N.H., Van Etten, J. and Goldstrohm, A.C. (2014) The RNA binding domain of Pumilio antagonizes poly-adenosine binding protein and accelerates deadenylation. RNA, 20, 1298–1319.

47. Go, C.D., Knight, J.D.R., Rajasekharan, A., Rathod, B., Hesketh, G.G., Abe, K.T., Youn, J.-Y., Samavarchi-Tehrani, P., Zhang, H., Zhu, L.Y., et al. (2021) A proximity-dependent biotinylation map of a human cell. Nature, 595, 120–124.

48. Gorgoni, B. and Gray, N.K. (2004) The roles of cytoplasmic poly(A)-binding proteins in regulating gene expression: a developmental perspective. Brief. Funct. Genomic. Proteomic., 3, 125–141.

49. Sachs, A.B., Davis, R.W. and Kornberg, R.D. (1987) A single domain of yeast poly(A)-binding protein is necessary and sufficient for RNA binding and cell viability. Mol. Cell. Biol., 7, 3268–3276.

50. Görlach, M., Burd, C.G. and Dreyfuss, G. (1994) The mRNA poly(A)-binding protein: localization, abundance, and RNA-binding specificity. Exp. Cell Res., 211, 400–407.

51. Kühn, U. and Pieler, T. (1996) Xenopus poly(A) binding protein: functional domains in RNA binding and protein-protein interaction. J. Mol. Biol., 256, 20–30.

52. Sawazaki, R., Imai, S., Yokogawa, M., Hosoda, N., Hoshino, S.-I., Mio, M., Mio, K., Shimada, I. and Osawa, M. (2018) Characterization of the multimeric structure of poly(A)-binding protein on a poly(A) tail. Sci. Rep., 8, 1455.

53. Xie, J., Kozlov, G. and Gehring, K. (2014) The ‘tale’ of poly(A) binding protein: the MLLE domain and PAM2-containing proteins. Biochim. Biophys. Acta, 1839, 1062–1068.

54. Fatscher, T., Boehm, V., Weiche, B. and Gehring, N.H. (2014) The interaction of cytoplasmic poly(A)-binding protein with eukaryotic initiation factor 4G suppresses nonsense-mediated mRNA decay. RNA, 20, 1579–1592.

55. He, S., Valkov, E., Cheloufi, S. and Murn, J. (2023) The nexus between RNA-binding proteins and their effectors. Nat. Rev. Genet., 24, 276–294.

56. Ludvigsen, M., Thorlacius-Ussing, L., Vorum, H., Moyer, M.P., Stender, M.T., Thorlacius-Ussing, O. and Honoré, B. (2020) Proteomic Characterization of Colorectal Cancer Cells versus Normal-Derived Colon Mucosa Cells: Approaching Identification of Novel Diagnostic Protein Biomarkers in Colorectal Cancer. Int. J. Mol. Sci., 21, 3466.

57. Nagaraj, N., Wisniewski, J.R., Geiger, T., Cox, J., Kircher, M., Kelso, J., Pääbo, S. and Mann, M. (2011) Deep proteome and transcriptome mapping of a human cancer cell line. Mol. Syst. Biol., 7, 548.

58. Zekri, L., Kuzuoğlu-Öztürk, D. and Izaurralde, E. (2013) GW182 proteins cause PABP dissociation from silenced miRNA targets in the absence of deadenylation. EMBO J., 32, 1052–1065.

59. Wolfe, M.B., Schagat, T.L., Paulsen, M.T., Magnuson, B., Ljungman, M., Park, D., Zhang, C., Campbell, Z.T., Goldstrohm, A.C. and Freddolino, P.L. (2020) Principles of mRNA control by human PUM proteins elucidated from multimodal experiments and integrative data analysis. RNA, 26, 1680–1703.

60. Li, S., Prasanna, X., Salo, V.T., Vattulainen, I. and Ikonen, E. (2019) An efficient auxin-inducible degron system with low basal degradation in human cells. Nat. Methods, 16, 866–869.

61. Sánchez, W.N., Driessen, A.J.M. and Wilson, C.A.M. (2025) Protein targeting to the ER membrane: multiple pathways and shared machinery. Crit. Rev. Biochem. Mol. Biol., 60, 33–79.

62. Deardorff, J.A. and Sachs, A.B. (1997) Differential effects of aromatic and charged residue substitutions in the RNA binding domains of the yeast poly(A)-binding protein. J. Mol. Biol., 269, 67–81.

63. Burd, C.G., Matunis, E.L. and Dreyfuss, G. (1991) The multiple RNA-binding domains of the mRNA poly(A)-binding protein have different RNA-binding activities. Mol. Cell. Biol., 11, 3419–3424.

64. Deo, R.C., Bonanno, J.B., Sonenberg, N. and Burley, S.K. (1999) Recognition of polyadenylate RNA by the poly(A)-binding protein. Cell, 98, 835–845.

65. Du, H., Zhao, Y., He, J., Zhang, Y., Xi, H., Liu, M., Ma, J. and Wu, L. (2016) YTHDF2 destabilizes m(6)A-containing RNA through direct recruitment of the CCR4-NOT deadenylase complex. Nat. Commun., 7, 12626.

66. Buschauer, R., Matsuo, Y., Sugiyama, T., Chen, Y.-H., Alhusaini, N., Sweet, T., Ikeuchi, K., Cheng, J., Matsuki, Y., Nobuta, R., et al. (2020) The Ccr4-Not complex monitors the translating ribosome for codon optimality. Science, 368, eaay6912.

67. Zhu, X., Cruz, V.E., Zhang, H., Erzberger, J.P. and Mendell, J.T. (2024) Specific tRNAs promote mRNA decay by recruiting the CCR4-NOT complex to translating ribosomes. Science, 386, eadq8587.

68. Höpfler, M., Absmeier, E., Peak-Chew, S.-Y., Vartholomaiou, E., Passmore, L.A., Gasic, I. and Hegde, R.S. (2023) Mechanism of ribosome-associated mRNA degradation during tubulin autoregulation. Mol. Cell, 83, 2290–2302.e13.

69. Chorghade, S., Seimetz, J., Emmons, R., Yang, J., Bresson, S.M., Lisio, M.D., Parise, G., Conrad, N.K. and Kalsotra, A. (2017) Poly(A) tail length regulates PABPC1 expression to tune translation in the heart. eLife, 6, e24139.

70. Yao, W., Yao, Y., He, W., Zhao, C., Liu, D., Wang, G. and Wang, Z. (2023) PABPC1 promotes cell proliferation and metastasis in pancreatic adenocarcinoma by regulating COL12A1 expression. Immun. Inflamm. Dis., 11, e919.

71. Pu, J., Zhang, T., Zhang, D., He, K., Chen, Y., Sun, X. and Long, W. (2021) High-Expression of Cytoplasmic Poly (A) Binding Protein 1 (PABPC1) as a Prognostic Biomarker for Early-Stage Esophageal Squamous Cell Carcinoma. Cancer Manag. Res., 13, 5361–5372.

72. Qi, Y., Wang, M. and Jiang, Q. (2022) PABPC1-mRNA stability, protein translation and tumorigenesis. Front. Oncol., 12, 1025291.

73. Bhandari, D., Raisch, T., Weichenrieder, O., Jonas, S. and Izaurralde, E. (2014) Structural basis for the Nanos-mediated recruitment of the CCR4-NOT complex and translational repression. Genes Dev., 28, 888–901.

74. Russell, P.J., Slivka, J.A., Boyle, E.P., Burghes, A.H.M. and Kearse, M.G. (2023) Translation reinitiation after uORFs does not fully protect mRNAs from nonsense-mediated decay. RNA, 6, 735–744.

75. Baluapuri, A., Zhao, N.C., Marina, R.J., Huang, K.-L., Kuzkina, A., Amodeo, M.E., Stein, C.B., Ahn, L.Y., Farr, J.S., Schaffer, A.E., et al. (2025) Integrator loss leads to dsRNA formation that triggers the integrated stress response. Cell, 188, 3184–3201.e21.

76. Lai, W.S., Arvola, R.M., Goldstrohm, A.C. and Blackshear, P.J. (2019) Inhibiting transcription in cultured metazoan cells with actinomycin D to monitor mRNA turnover. Methods, 155, 77–87.

77. Arvola, R.M. and Goldstrohm, A.C. (2024) Measuring Poly-Adenosine Tail Length of RNAs by High-Resolution Northern Blotting Coupled with RNase H Cleavage. Methods Mol. Biol. Clifton NJ, 2723, 93–111.

78. McKenney, K.M., Connacher, R.P., Dunshee, E.B. and Goldstrohm, A.C. (2024) Chemi- Northern: a versatile chemiluminescent northern blot method for analysis and quantitation of RNA molecules. RNA, 30, 448–462.

