## Supplemental Figures for "Cytoplasmic poly-adenosine binding proteins modulate susceptibility of mRNAs to RNA-binding protein-directed decay"

**SUPPORTING INFORMATION**
**McKenney et al.**

**Supplemental Figure S1. Comparison of Nluc luciferase expression from reporter mRNAs shown in Figure 1.**
The relative response ratio (RRR) for the indicated reporters in HCT116 cells are graphed. The RRR values were calculated as the ratio of Nluc/Fluc activity, as described in Materials and Methods. n=9; 3 experiments, each with 3 biological replicates +/- SD.

**Supplemental Figure S1**
**McKenney et al.**

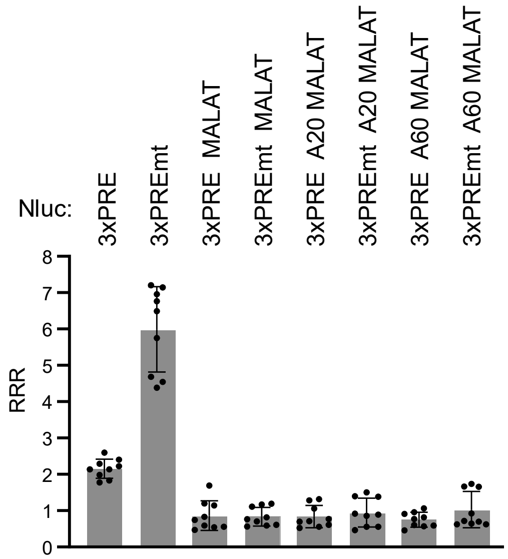

**Supplemental Figure S2. Depletion of PABPC1 reduces PUM-mediated repression.**

**A.** Western blot analysis confirms RNAi-mediated depletion of PABPC1 protein in samples from a representative experimental replicate of reporter assays shown in panel B. Western blot of GAPDH served as a loading control. PUM1 and PUM2 levels were assessed by western blot.

**B.** Luciferase reporter assay showing the effect of PABPC1 knockdown on PUM repression of the Nluc 3xPRE reporter, relative to the mutant version, Nluc 3xPREmt. PABPC1 was depleted by RNAi and compared to non-targeting siRNA control (NTC). n=9; 3 experiments, each with 3 biological replicates; +/- SD.

**C.** Western blot of PABPC1 depletion using the Auxin Inducible Degron (AID) in samples from a representative experimental replicate of the experiment in Panel D. GAPDH served as a loading control.

**D**. Luciferase reporter assay showing the effect of PABPC1-AID depletion on PUM repression of the 3xPRE reporter, relative to the mutant reporter, Nluc 3xPREmt. PABPC1-AID was degraded upon treatment with the auxin, indole-3-acetic acid (+IAA), and compared to a vehicle-only control (-IAA). n=9; 3 experiments, each with 3 biological replicates; +/- SD.

**E**. Expression of the Firefly luciferase MALAT1 (Fluc MALAT1) internal control was not substantially affected by PABPC1 and PABPC4 depletion. The Fluc MALAT1 activity was measured in relative light units (RLU) from reporter assays under PABPC1&4 depletion conditions as shown in Figure 3A, 3B.

**F**. Fluc MALAT1 expression, measured as Firefly luciferase activity (RLUs), was not substantially changed by over-expression of Halotag-PABPC1 fusion. The data in the graph is from the experiment shown in Figure 7A, 7B. n=9; 3 experiments, each with 3 biological replicates; +/- SD.

**Supplemental Figure S2**
**McKenney et al.**

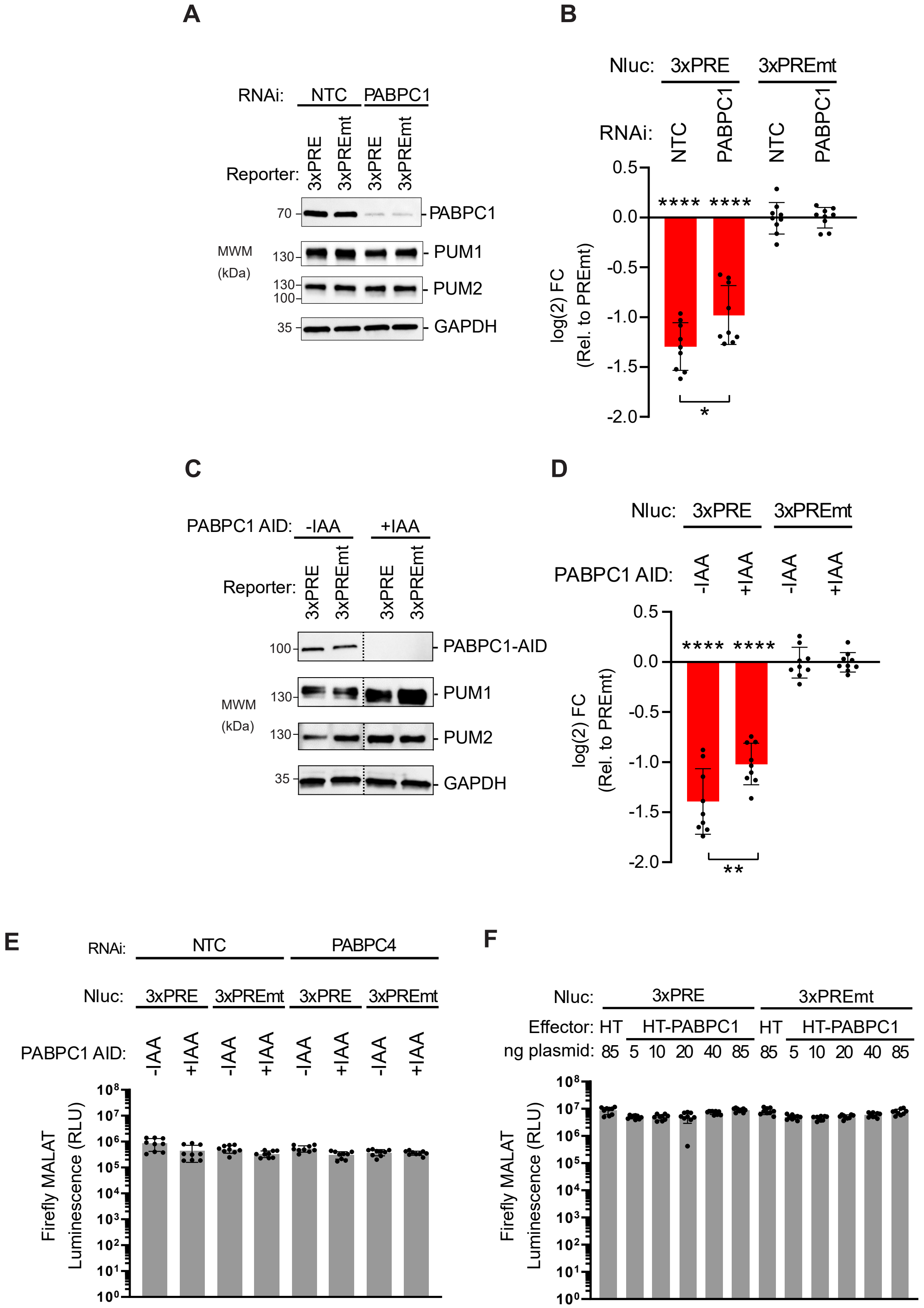

**Supplemental Figure S3. PABPC1&4 depletion destabilizes mRNAs.**

**A.** The Nluc 6xPRE reporter is robustly repressed in HCT116 cells relative to the mutant version, Nluc 6xPREmt. Depletion of both PABPC1 and PABPC4 alleviated PUM repression of the Nluc 6xPRE. Depletion of PABPC1-AID was induced by addition of IAA, compared to vehicle (-IAA). Depletion of PABPC4 was achieved by RNAi using specific siRNAs, compared to the nontargeting control siRNA (NTC). n=9; 3 experiments, each with 3 biological replicates; +/- SD. For significance calling, $p < 0.05$ = *, $p < 0.01$ = **, $p < 0.001$ = ***, $p < 0.0001$ = **** based on ordinary one-way ANOVA and Tukey test for multiple comparisons. Significant differences indicated above the X-axis are relative to the Nluc 6xPREmt reporter.

**B.** Products from PCR-based genotyping of the PABPC1 AID in HCT116 AtAFB2 cells compared to a wild type control sample were analyzed by ethidium bromide stained agarose gel electrophoresis. The larger 1.2kb PABPC1 AID PCR product demonstrates homozygous tagging of the PABPC1 locus, relative to the 0.9kb PCR product in the parental wild type (WT) HCT116 AtAFB2 cell line.

**C.** Western blot confirmation of depletion of PABPC1 AID after 3 hours treatment with indole-3-acetic acid (+IAA). A shift in molecular weight of PABPC AID (83 kDa) is observed compared to untagged PABPC1 (71 kDa) in the wild type HCT116 control. GAPDH served as loading control.

**D.** Northern blots of Nluc reporters and the 18S ribosomal rRNA from the 2 additional experimental replicates from the Tet-off transcription shut-off experiment shown in Figure 5.

**E.** Corresponding western blot verification of PABPC1 AID and PABPC4 RNAi depletion for experiment replicates shown in Panel D.

**F.** Ethidium bromide stained, denaturing formaldehyde agarose gels of the RNA samples from the northern blots of the 3 experimental replicates shown in Panel D and Figure 5. Visualization of the 28S and 18S rRNA verifies equal loading and integrity of the RNA samples.

**Supplemental Figure S3**
**McKenney et al.**

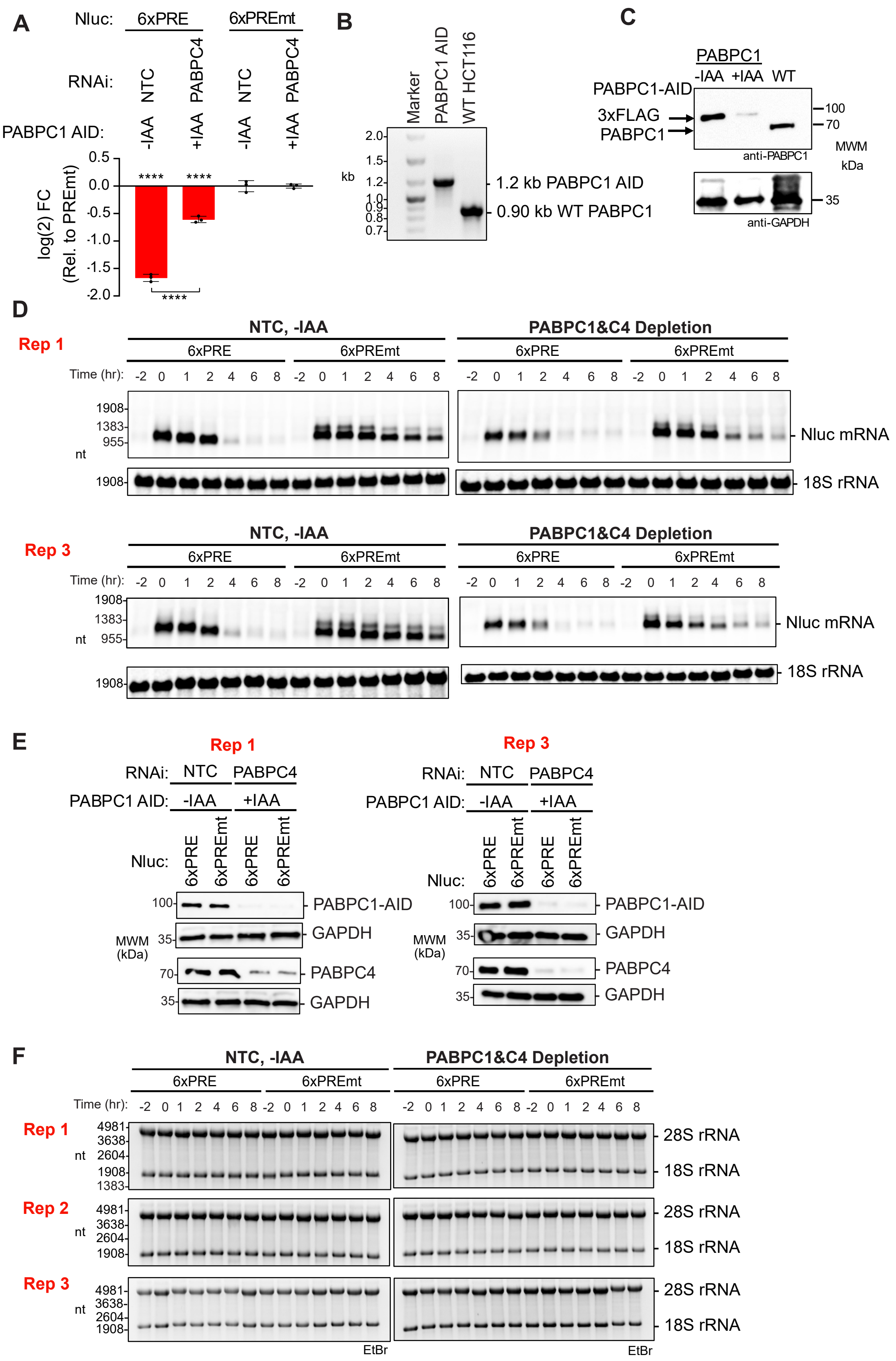

**Supplemental Figure S4. over-expression of cytoplasmic PABP homologs alleviates PUM-repression.**

**A.** Western blot analysis of titration of transfected, untagged PABPC1 from a representative experimental replicate for Panel B. Cells transfected with Halotag (HT) served as a control. GAPDH served as a loading control. PUM1 and PUM2 protein expression was assessed as a control. Below is a graph of the fold increase in PABPC1 expression over the level of endogenous PABPC1 in the cells that were transfected with the HT control. n=3 experimental replicates; +/- SD.

**B.** Reporter assay showing effect of titration of transfected, untagged PABPC1 on PUM repression of Nluc 3xPRE reporter, relative to mutant control reporter, in wild type HCT116 cells. n=9; 3 experiments, each with 3 biological replicates; +/- SD.

**C.** Diagram of the domain architecture for four human PABPC paralogs tested in this study.

**D.** Western blot analysis of titration of untagged PABPC4 from a representative experimental replicate of the results shown in Panel E. GAPDH served as a loading control. PUM1 and PUM2 protein expression was measured as a control.

**E.** Reporter assay showing effect of PABPC4 over-expression on PUM repression of Nluc 3xPRE reporter, relative to mutant control, in wild type HCT116 cells. n=9; 3 experiments, each with 3 biological replicates; +/- SD.

**F.** Western blot analysis of PABPC paralogs, including PABPC1, PABPC3, PABPC4, and PABPC5, at 85 ng transfected plasmid, from a representative experimental replicate of results shown in Panel G. GAPDH or H3 served as loading controls.

**G.** Reporter assay showing effect over-expression of PABPC paralogs on PUM repression of Nluc 3xPRE reporter, relative to mutant version, in wild type HCT116 cells. n=9; 3 experiments, each with 3 biological replicates; +/- SD. For significance calling, $p < 0.05$ = *, $p < 0.01$ = **, $p < 0.001$ = ***, $p < 0.0001$ = **** based on ordinary one-way ANOVA and Tukey test for multiple comparisons.

### Supplemental Figure S4
McKenney et al.

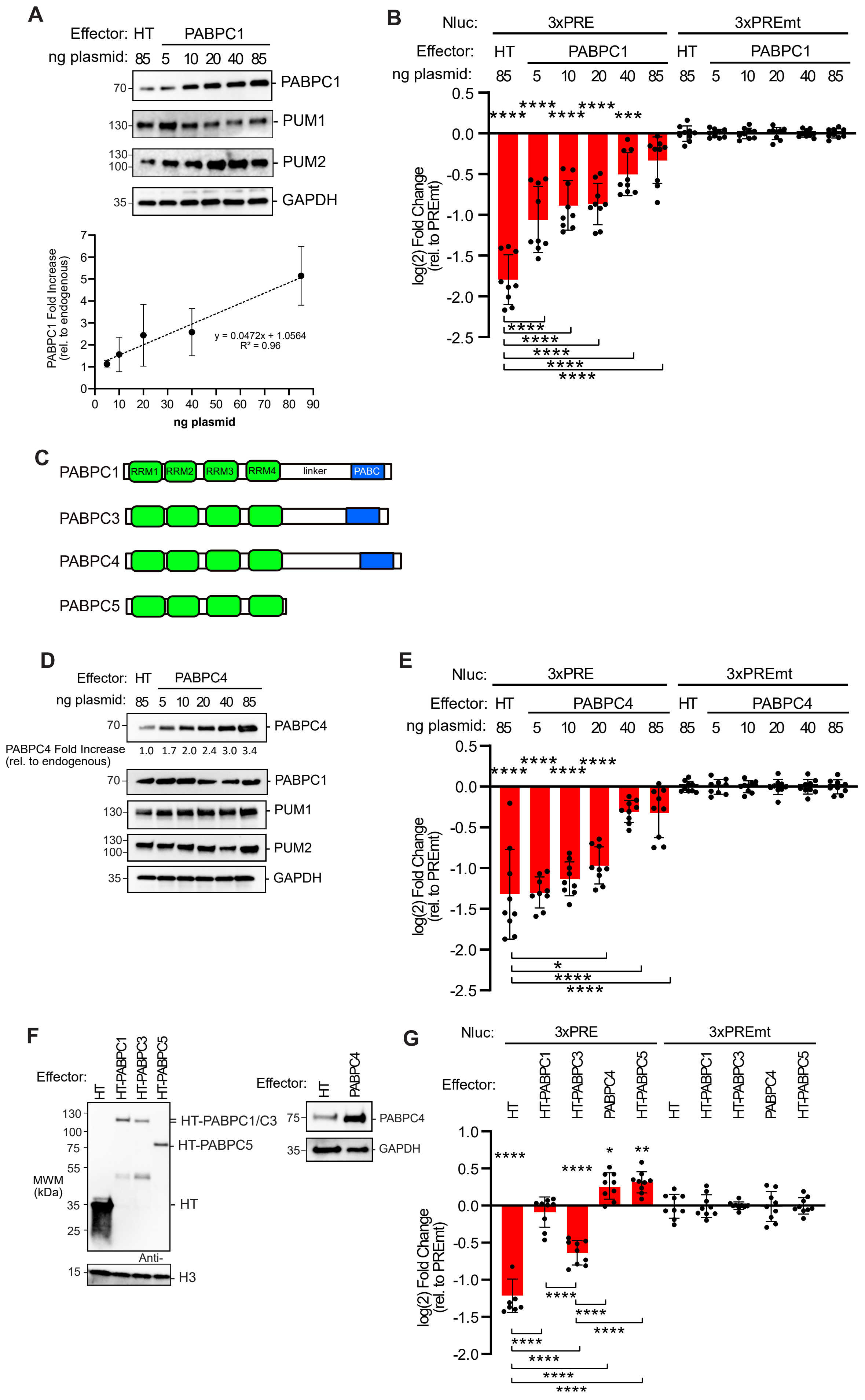

**Supplemental Figure S5. PABPC1 over-expression stabilizes mRNAs.**

**A**. Northern blots of Nluc reporters and the 18S ribosomal rRNA from the 2 additional experimental replicates (Rep 2 and Rep3) from the Tet-off transcription shut-off experiment with over-expressed HT-PABPC1 shown in Figure 9E.

**B**. Corresponding western blot verification of HT-PABPC1 over-expression for the two additional experiment replicates shown in Panel A. Both over-expressed HT-PABPC1 and endogenous PABPC1 were detected with a PABPC1 antibody. HT was detected in the control samples. Histone H3 served as a loading control.

**C**. Ethidium bromide stained, denaturing formaldehyde agarose gels of the RNA samples from the northern blots of the 3 experimental replicates shown in Panel A and Figure 9E. Visualization of the 28S and 18S rRNA verifies equal loading and integrity of the RNA samples.

### Supplemental Figure S5
McKenney et al.

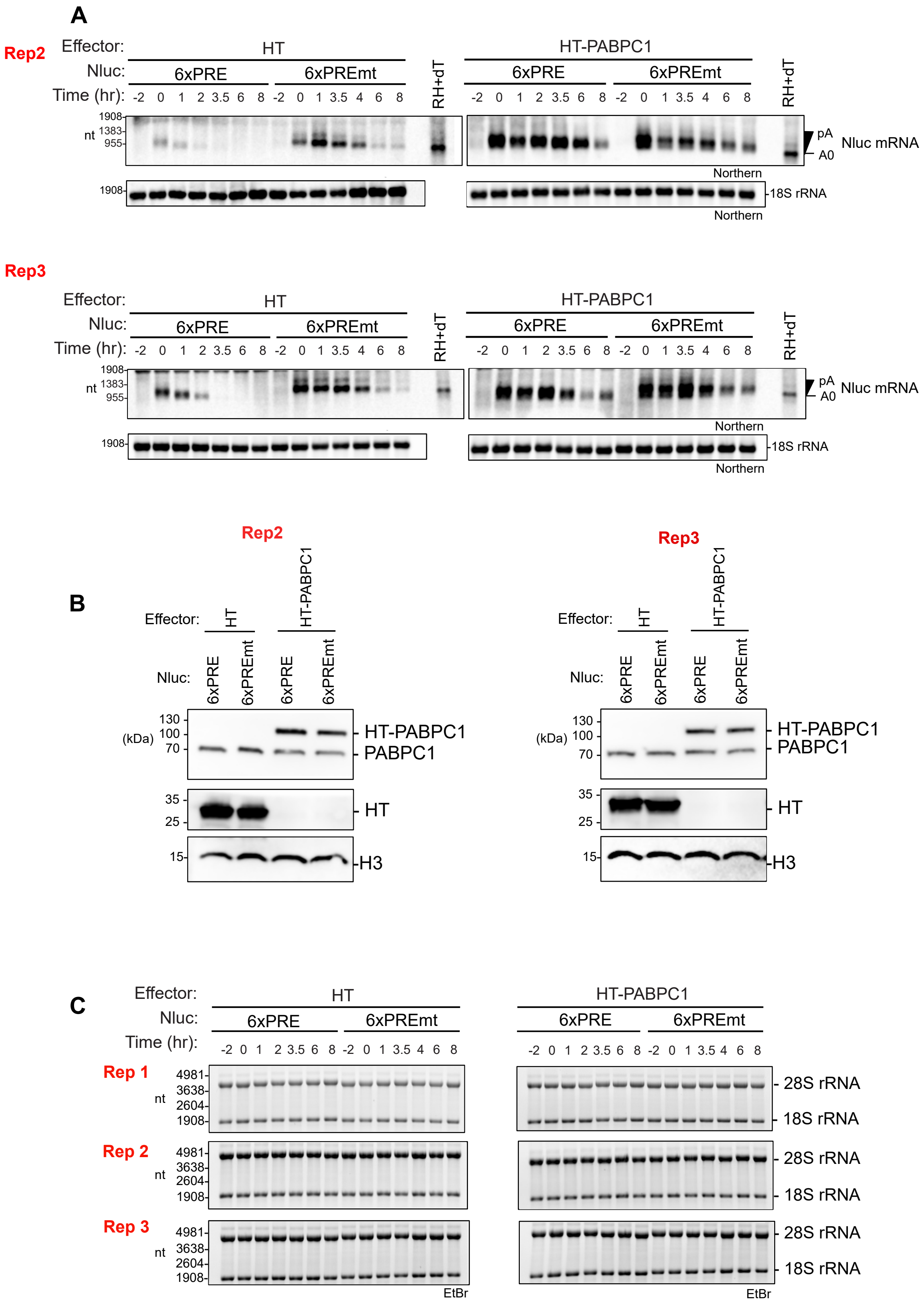

**Supplemental Figure S6. PABPC1&4 mRNA expression levels across normal human tissues.**

Violin plots of PABPC1 and PABPC4 mRNA levels reported as normalized expression units (transcripts per million reads, TPM). This RNA-seq data was obtained from the Genotype-Tissue Expression (GTEX) database for normal human tissues (version 10, Gencode version 39, STAR v2.7.10a, transcript quantification: RSEM v1.3.3) and included 54 tissues from 946 donors and 19788 samples. Plots including distributions of TPM measurements along with mean and quartile values. Tissues are ranked from highest to lowest, left to right, respectively. Ranges of the mean values are indicated on the right.

**Supplemental Figure S6**
**McKenney et al.**

A

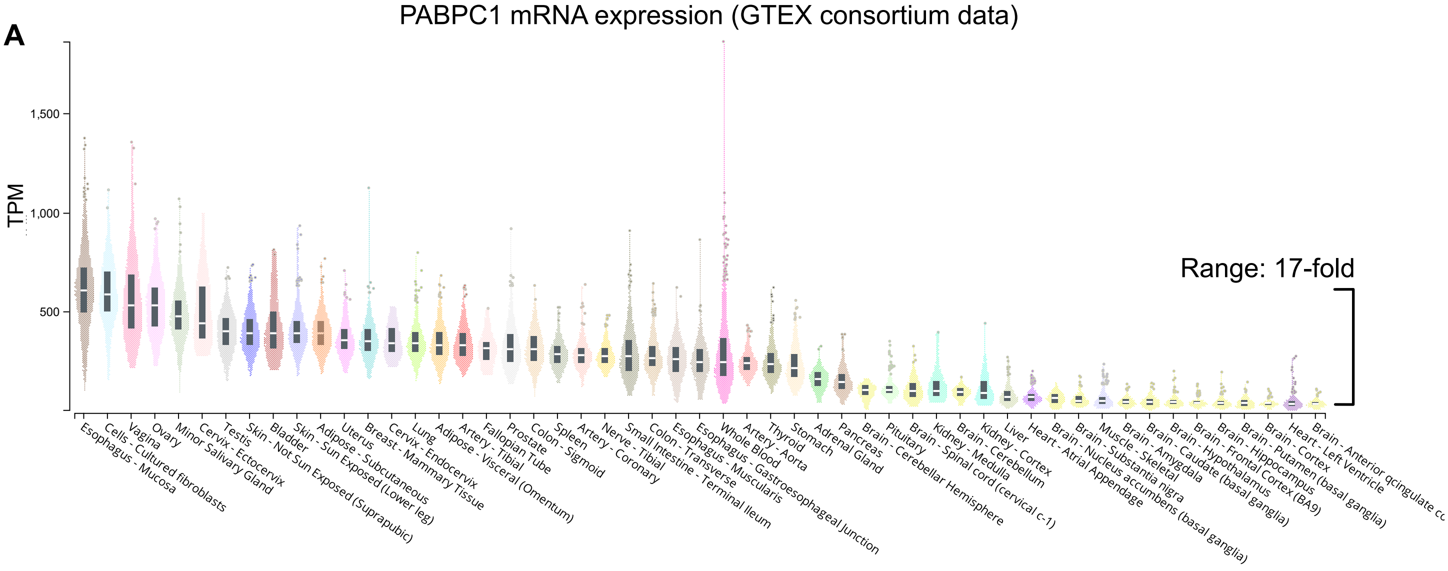


B

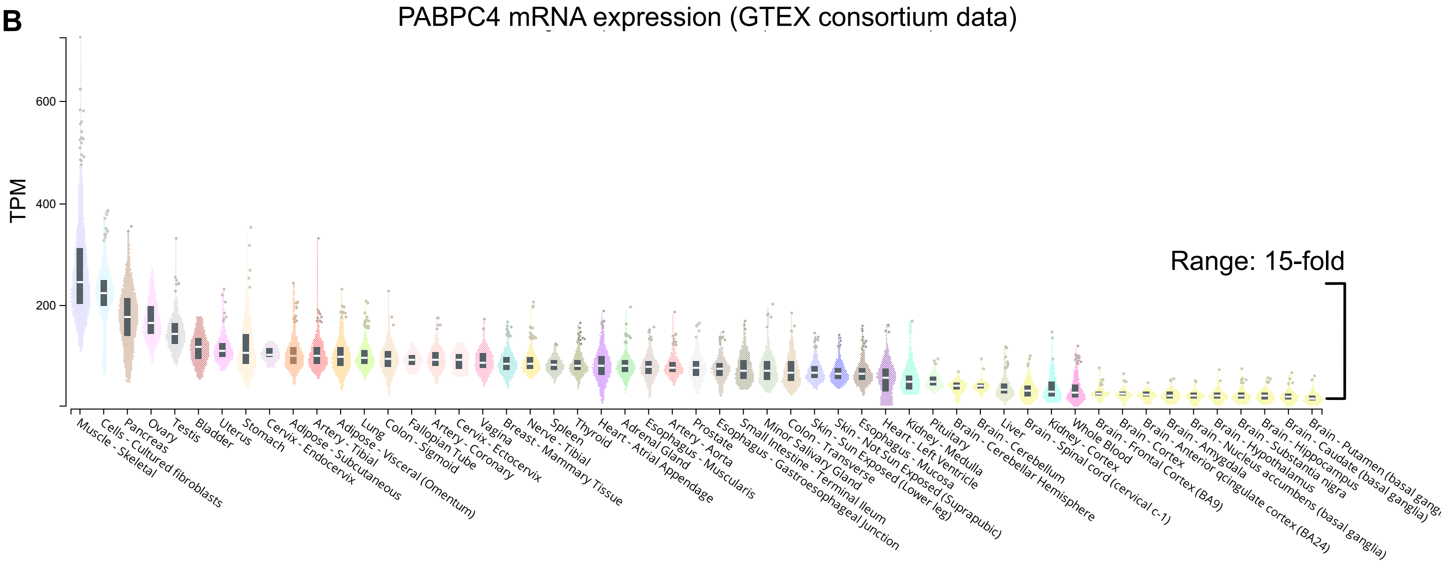

**Supplemental Figure S7. PABPC protein expression across normal human tissues.**
Analysis of protein expression of PABPC in human tissues from donors with no known disease. PABPC1, PABPC3, PABPC4, and GAPDH proteins were detected on a human tissue western blots (INSTA-Blot̉ Human Tissues, NBP2-30113) with 20 μg per lane of human tissue lysates from brain, heart, small intestine, kidney, liver, lung, skeletal muscle, stomach, spleen, ovary, and testis, as indicated at the top. Each blot included a molecular weight marker (MWM, Thermo Scientific, LC5677). Total protein in each lane was detected by staining the PVDF membrane with amido black. Detected bands consistent with the expected molecular weight of each target protein are indicated on the right side of each blot.
**A.** Western blot with an antibody specific for PABPC1 (Abcam, ab6125).
**B**. Western blot with antibody that recognizes PABPC1 along with PABPC3 and PABPC4 (Cell Signaling Technologies, 4992S).
**C.** Western blot with an antibody that detects PABPC4 (Bethyl labs, A301-467A). This blot was probed with GAPDH antibody as a loading control (bottom panel).

**Supplemental Figure S7**
**McKenney et al.**

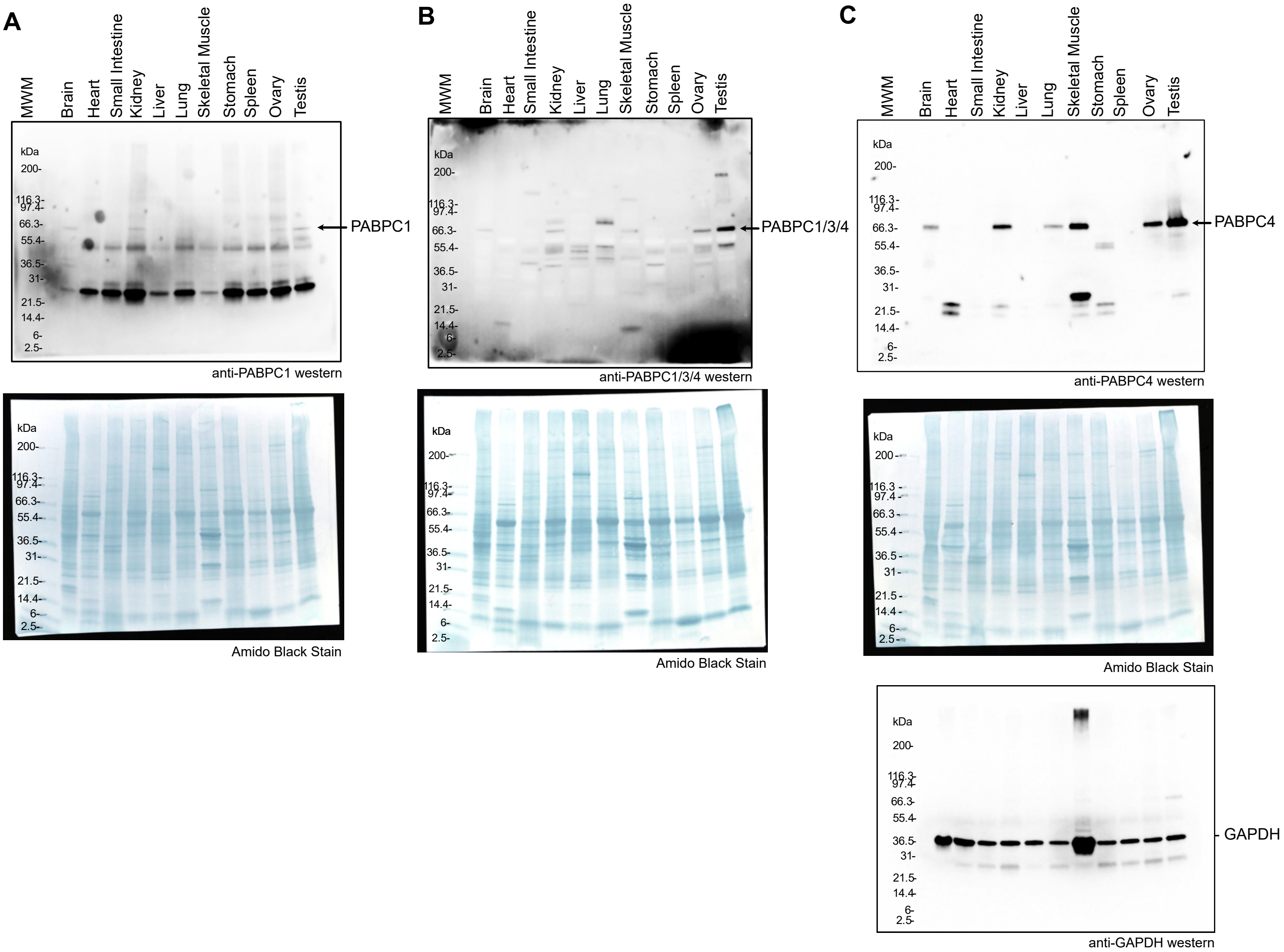
